# A faithful in vivo model of human macrophages in metastatic melanoma

**DOI:** 10.1101/2021.09.09.459682

**Authors:** Valentin Voillet, Trisha R. Berger, Kelly M. McKenna, Kelly G. Paulson, Kimberly S. Smythe, Daniel S. Hunter, William J. Valente, Stephanie Weaver, Jean S. Campbell, Teresa S. Kim, David R. Byrd, Jason H. Bielas, Robert H. Pierce, Aude G. Chapuis, Raphaël Gottardo, Anthony Rongvaux

**Author notes:** **Corresponding author:** Anthony Rongvaux; Fred Hutchinson Cancer Research Center, 1100 Fairview Ave. N. – D3-100, Seattle, WA 98109 USA. These authors contributed equally (Valentin Voillet, Trisha R. Berger).

## Abstract

Despite recent therapeutic progress, advanced melanoma remains lethal for many patients. The composition of the immune tumor microenvironment (TME) has decisive impacts on therapy response and disease outcome. High dimensional analyses of patient samples can reveal the composition and heterogeneity of the immune TME. In particular, macrophages are known for their cancer-supportive role, but the underlying mechanisms are incompletely understood, and experimental in vivo systems are needed to test the functional properties of these cells. We characterized a humanized mouse model, reconstituted with a human immune system and a human melanoma, in which: (1) human macrophages support metastatic spread of the tumor; and (2) tumor-infiltrating macrophages have a specific transcriptional signature that faithfully represents the transcriptome of macrophages from patient melanoma samples and is associated with shorter survival. This model complements patient sample analyses, enabling the elucidation of fundamental principles in melanoma biology, and the development and evaluation of candidate therapies.

## Introduction

Almost 60,000 people die annually of melanoma worldwide [1]. Recent therapeutic progress has profoundly transformed clinical outcomes in advanced melanoma patients. With the introduction of oncogene-targeted and immune checkpoint-blocking therapies, the 5-year survival rate has increased to over 50% [2, 3]. For patients whose tumors are resistant to available therapies, metastasis is the most frequent cause of death [4, 5]. Indeed, data from 2009 indicated that the five-year survival rate for patients with non-metastatic melanoma ranged between 53% and 97%, but was less than 20% for patients with metastatic disease [6]. Therefore, new therapeutic strategies targeting the mechanisms underlying metastatic spread are needed to further improve clinical outcomes [3]. To reach this goal, predictive experimental models are needed to extend our understanding of the fundamental biology of metastatic melanoma, and for pre-clinical evaluations of candidate drugs [7].

The immune composition of the tumor microenvironment (TME) has decisive impacts on disease progression and therapy responsiveness [8]. In particular, a high density of TME-infiltrating macrophages is associated with unfavorable clinical outcomes in melanoma and most other cancer types [9, 10]. Regarding how macrophages support cancer progression, they promote tumor vascularization, suppress adaptive immunity against tumor antigens, and favor the metastatic spread of the tumor [11]. Macrophages are heterogeneous and multifunctional cells of the mononuclear phagocyte system (MPS), which also includes monocytes and dendritic cells [12]. They are well known for their role in host defense against infections, but they also play a variety of functions in tissue homeostasis in both healthy and pathological conditions [13]. In the past few years, cutting-edge technologies, such as *in vivo* fate-mapping, epigenomics and single cell genomics, have elucidated the ontogeny and diversity of macrophages and other MPS cells in the mouse [13, 14]. However, mice are not humans, and major differences exist between MPS cells of these two species [15]. In humans, research on macrophages and their physiological microenvironment is frequently limited to observational studies. For example, a recent systemic analysis of 15 human cancer tumors generated an atlas of tumor-infiltrating MPS cells [16]. This rich resource provides a systemic view of the heterogeneity of MPS cells in human cancer. But, apart from clinical trials, functional *in vivo* studies of these cells represent a considerable challenge. Consequently, tumor-associated macrophages remain a largely untapped target for the treatment of cancer [17].

Genetically engineered mouse (GEM) models of melanoma, such as mice with inducible BRAF and PTEN mutations, recapitulate recurrent tumor-driving mutations found in human melanoma, and enable studies of cancer initiation and progression in immunocompetent mice [18]. Such models are invaluable research tools but do not fully represent the biology of human melanoma or human MPS cells [7]. In patient-derived xenografts (PDX) or cell line-derived xenografts (CDX), a human tumor is implanted into an immunodeficient recipient mouse [19]. PDXs can capture the heterogeneity of human melanoma as well as their responsiveness to candidate drugs [20]. However, as the mouse host lacks most lymphoid cells, and because the remaining mouse immune cells are not fully cross-reactive with the human tumor, standard PDX and CDX models have limited applicability for studies of human cancer-immune cell interactions [19]. This limitation can be overcome with “immuno-PDX” or “immuno-CDX” models, in which the recipient mouse is repopulated with a humanized immune system [7, 19, 21]. With the advent of a new-generation of “humanized mice” that support the development and function of human myeloid cells [22, 23], models recapitulating the tumor-supportive role of human macrophages in human melanoma are becoming available.

Here, we characterize an immuno-CDX model of melanoma and demonstrate that metastatic spread of the human tumor is supported by human MPS cells. Through single-cell transcriptomic profiling, we describe a characteristic, TME-associated MPS cell population. Similar cells are present in patient melanoma samples and are associated with unfavorable outcomes. These results open new opportunities for the development and clinically-relevant *in vivo* evaluation of therapies specifically targeting the metastasis-supportive activities of human macrophages in melanoma.

## Results

### Macrophages support metastasis in a humanized mouse model of melanoma

MISTRG (M-CSF^h/h^ IL-3/GM-CSF^h/h^ SIRPα^h/m^ TPO^h/h^ RAG2^-/-^ IL-2Rγ^-/-^) is an immunodeficient mouse strain in which several cytokine-encoding genes are humanized by knock-in replacement of the mouse alleles with their human counterparts, from the start to the stop codon [22-24]. Upon transplantation of human CD34^+^ hematopoietic stem and progenitor cells (HSPCs) into preconditioned newborn MISTRG mice, following a standard protocol [22], a multi-lineage human immune system develops, including B and T lymphocytes, natural killer (NK) cells, monocytes, macrophages and dendritic cells (DCs) [22, 23]. To specifically study human M-CSF-dependent MPS cells in this humanized mouse system, we also generated ISTRG mice (IL-3/GM-CSF^h/h^ SIRPα^h/m^ TPO^h/h^ RAG2^-/-^ IL-2Rγ^-/-^) that lack the humanized M-CSF-encoding allele. MISTRG and ISTRG support high levels of human CD45^+^ immune cell chimerism, reaching 60-80% in the blood 16 weeks post-transplantation of fetal CD34^+^ HSPCs (**Figure 1A**). However, unlike MISTRG, ISTRG mice support limited development of blood CD33^+^ myeloid cells (**Figure 1B**) and of macrophages in tissues (identified with a CD68/CD163 antibody cocktail, **Figure 1C**). When we implant the human Me275 melanoma cell line [25] subcutaneously in the flank of these humanized mice, CD68^+^/CD163^+^ macrophages robustly infiltrate the tumors in MISTRG, but less so in ISTRG mice (**Figure 1D**). The differential abundance of MPS cells in MISTRG versus ISTRG does not affect the growth of primary tumors (**Figure 1E**). However, the presence of human MPS cells has a drastic effect on tumor metastasis. Indeed, we observe higher counts of metastatic nodules in the liver and spleen of humanized MISTRG mice than in humanized ISTRG mice (**Figure 1F**). We confirmed this observation by histology, staining humanized mouse tissues with an antibody cocktail specific for melanoma antigens (**Figure 1G**) and enumerating melanoma cells in liver and lung (**Figure 1H**). These observations show that human macrophages support cancer progression, particularly metastatic spread. Therefore, the humanized MISTRG mouse model of human melanoma recapitulates observations in patients and from regular mouse models.

**Figure 1.**
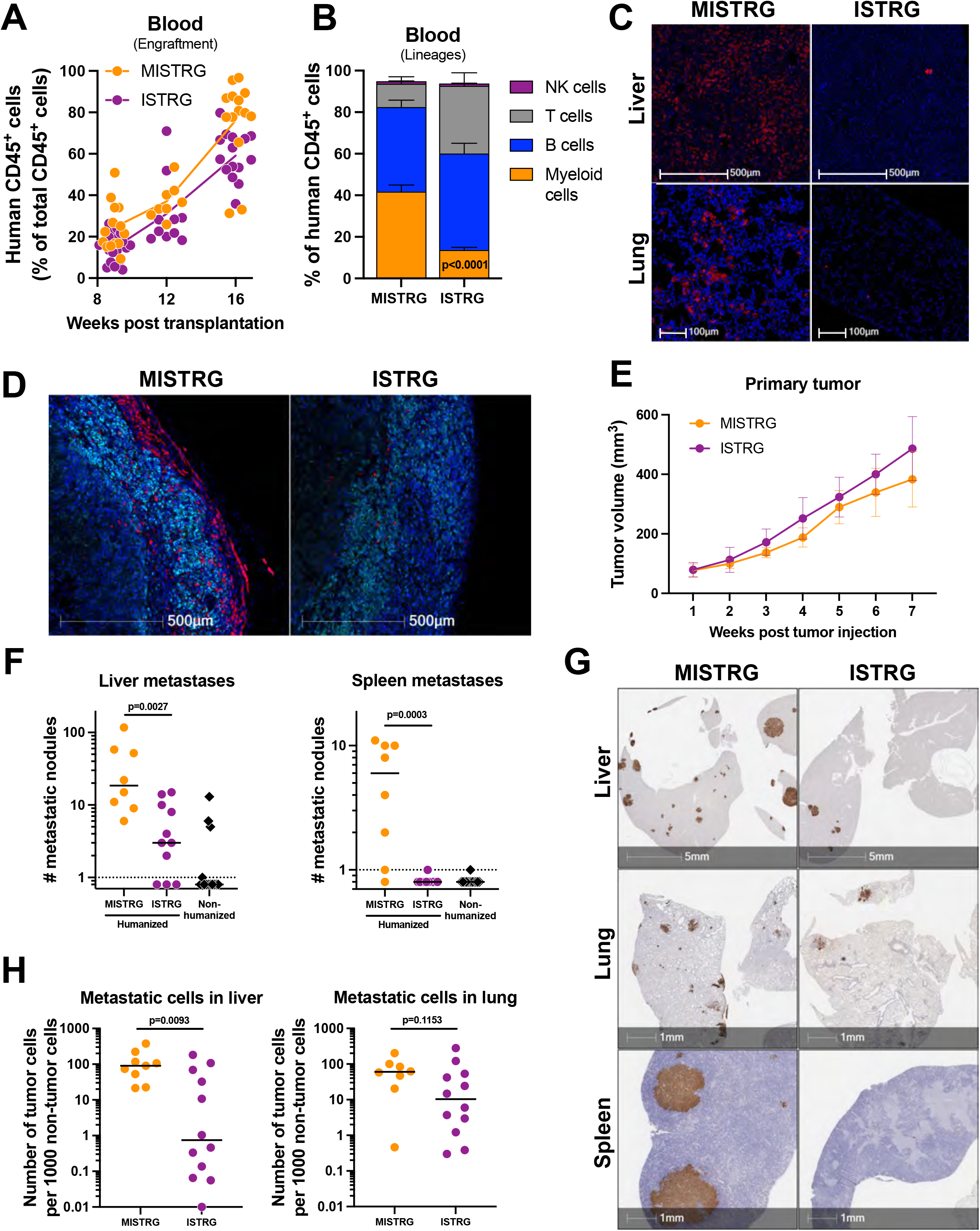
Macrophages support metastatic spread in a humanized mouse melanoma model. (**A** and **B**) Pre-conditioned MISTRG and ISTRG mice were humanized by transplantation of fetal CD34^+^ cells. Overall chimerism level, measured by flow cytometry as the percentage of human CD45^+^ cells among total CD45^+^ cells in the blood, was determined 9, 12 and 16 weeks post-transplantation (n=14(18) MISTRG(ISTRG), each symbol represents an individual mouse) (**A**). Human immune cell lineage composition was determined 12 weeks post-transplantation (n=18(18) MISTRG(ISTRG), error bars indicate mean ± S.E.M, p value calculated by the Mann–Whitney test) (**B**). (**C**) Human tissue-infiltrating macrophages were identified by immuno-histochemistry (IHC) using an anti-macrophage (CD68/CD163) antibody cocktail (red), with DAPI counterstaining of nuclei (blue). Representative images of 6 mice of each genotype, transplanted with human CD34^+^ cells from two individual donors. (**D**) Me275 human melanoma cells were implanted subcutaneously in the flank of humanized MISTRG and ISTRG mice. Three weeks later, tumor-infiltrating macrophages were identified by IHC using an anti-CD68/CD163 antibody cocktail (red). Tumor cells were stained with an anti-melanoma antibody cocktail (cyan) and nuclei counterstained with DAPI (blue). (**E** and **F**) Me275 tumor size was monitored weekly (n=8(11) MISTRG(ISTRG), error bars indicate mean ± S.D.) (**E**). Metastatic nodules were enumerated macroscopically on liver and spleen after necropsy, 7 weeks post tumor implantation (n=8(11) MISTRG(ISTRG), p values calculated by the Mann–Whitney test) (**F**). (**G** and **H**) Melanoma cells were identified by IHC in the indicated tissues, using an anti-melanoma antibody cocktail (brown) (**G**). The frequency of tumor cells, among total nucleated cells, were quantified using the HALO software (n=9(12) MISTRG(ISTRG), p values calculated by the Mann–Whitney test) (**H**).

### A distinct MPS cell population infiltrates melanoma primary tumor

To study the composition of the immune TME, we isolated human CD45^+^ immune cells from Me275 melanoma implanted in MISTRG mice, as well as from control tissues (bone marrow, liver and lung) of non-implanted mice. We analyzed these cells by single-cell RNA sequencing (scRNAseq), using the 10x Genomics platform [26]. We visualized all cells from all tissues (n = 6,399, **Suppl. Fig 1A**) with Uniform Manifold Approximation and Projection (UMAP). Unsupervised clustering distinguished diverse human immune cell lineages, including bone marrow (BM) progenitors, erythroid cells, B and T lymphocytes, NK cells and MPS cells, among human CD45^+^ cells from MISTRG (**Figure 2A, Suppl. Fig 1B** and **Suppl. Table 1A**). Some cell clusters were present almost exclusively in a specific tissue, such as BM progenitors (clusters 8, 13, 17 and 18) or liver MPS cells (cluster 12). In contrast, most clusters were distributed across tissues (**Figure 2B**). These observations suggest that cell clustering is driven by the biology of the cells, and not by technical biases introduced during cell isolation from different tissues. In the tumor, MPS cells were the most abundant cells (63.3%, including 7.5% CLEC9A^+^ DCs and 6.3% plasmacytoid DCs), followed by T cells (28.5%, including 10.5% FOXP3^+^ regulatory T cells), NK cells (6.1%) and B cells (2.1%) (**Figure 2C, Suppl. Fig 1C** and **Suppl. Table 1B**). This human immune TME composition was confirmed by multiplexed immunohistochemistry (mIHC) analysis, showing abundant and widespread CD163^+^ MPS cells, localized CD3^+^ T cells and rare CLEC9A^+^ DCs (**Figure 2D**).

**Figure 2.**
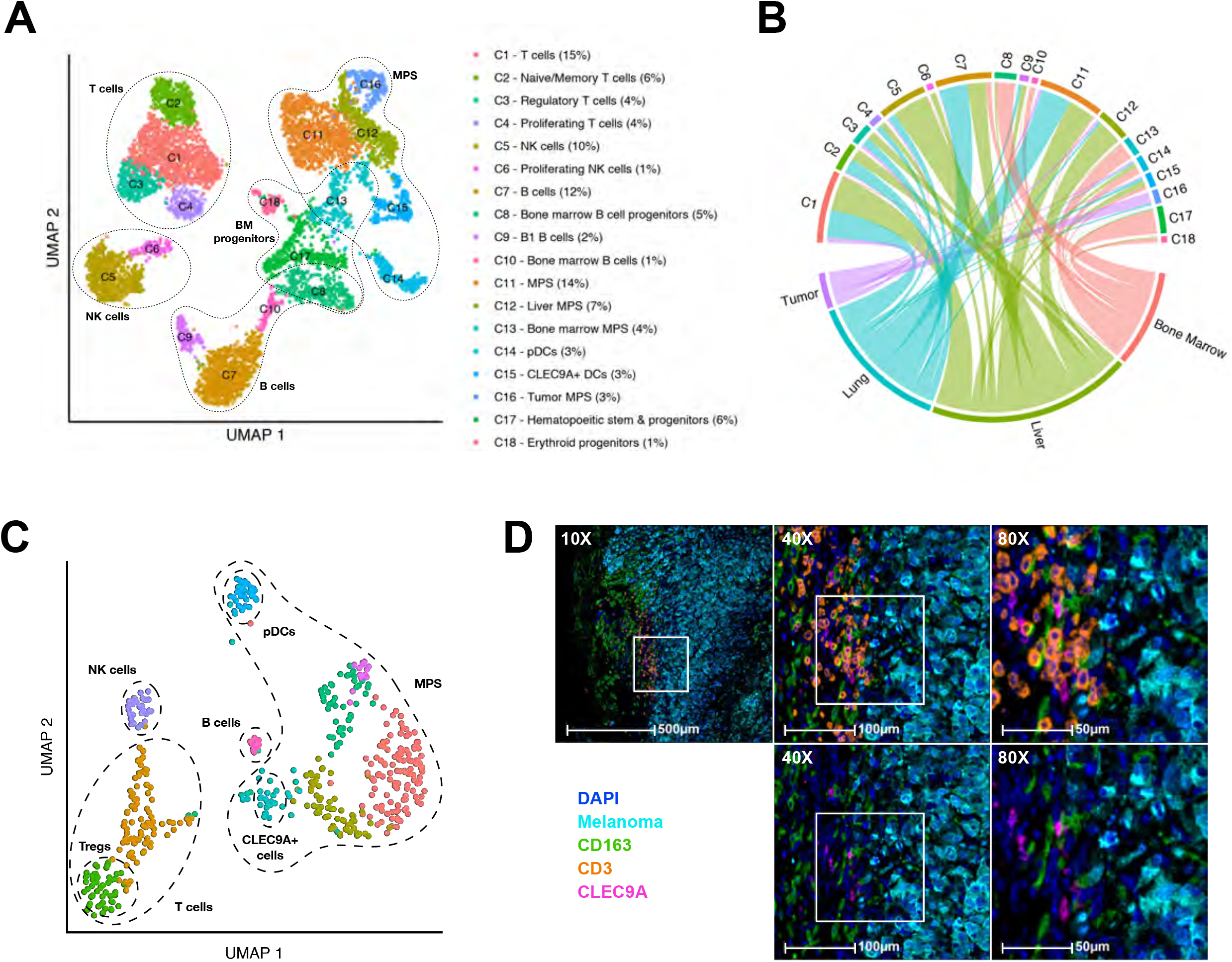
scRNAseq analysis of human immune cells in MISTRG tissues and melanoma. (**A**) UMAP embedding of all human CD45^+^ cells (n = 6,399 cells; Suppl. Fig 1A) isolated from tissues (bone marrow, liver and lung) of naïve humanized MISTRG mice, and from the tumor microenvironment of Me275 melanoma-bearing humanized MISTRG. Cell clusters are annotated based on expression of the representative marker genes shown in Suppl. Fig. 1B and Suppl. Table 1A. (**B**) Circular plot illustrating the distribution of clusters C1-C18 across tissues. (**C**) UMAP plot of human CD45^+^ cells from the Me275 tumor microenvironment colored by clusters. Representative marker genes used to annotate each cell cluster are shown in Suppl. Fig 1D and Suppl. Table 1A. (**D**) mIHC analysis of human immune cells (CD163^+^ macrophages, CD3^+^ T cells and CLEC9A^+^ DCs) in the microenvironment of Me275 melanoma.

Since human MPS cells supported tumor progression in MISTRG (**Figure 1**), we computationally isolated these cells (2,152 cells, **Suppl. Fig 2A**) and re-analyzed them at higher resolution. We identified a total of eleven MPS cell clusters across all tissues (**Figure 3A, B**). Among MPS cells, DC subsets are easily identifiable, based on the expression of defining markers (**Suppl. Fig 2B** and **Suppl. Table 2A**): CD1c^+^ DCs (cDC2) in the MPS5 cluster; CLEC9A^+^ DCs (cDC1) in MPS6 and plasmacytoid dendritic cells (pDCs) in MPS9 and MPS10. Finally, the MPS11 cluster corresponds to the recently identified LAMP3^+^ DC subset in the humanized mice [27]. Together, these results demonstrate that the MISTRG model faithfully recapitulates the diversity of well-described human DC subsets. In contrast to DC clusters that are distributed across tissues, other MPS cell clusters (i.e., monocytes or macrophages) displayed a more tissue-specific distribution (**Figure 3B**): MPS7 and MPS8 clusters correspond to BM granulocytic and monocytic progenitors; MPS1 was present mostly in the liver; and MPS2 and MPS3 were predominantly found in the lung. This observation is in line with results showing that the epigenomic landscape and transcriptional profile of macrophages is instructed by tissue environment cues in mice [28, 29]. Of interest, MPS4 cells were present almost exclusively in tumors, represented over 75% of intratumoral MPS cells (**Figure 3B**), and presented a clearly distinct transcriptional signature when the most differentially expressed genes of each cluster were compared (**Figure 3C** and **Suppl. Table 2A**). Within the transcriptional signature of MPS4 cells, we selected four highly overexpressed genes (**Figure 3D**) that encode proteins (CD14, CD163, MARCO and CTSL) for which antibodies are commercially available and validated for IHC. We found that these protein markers were co-expressed by most CD14^+^ MPS4 cells found in the Me275 TME (**Figure 3E**).

**Figure 3.**
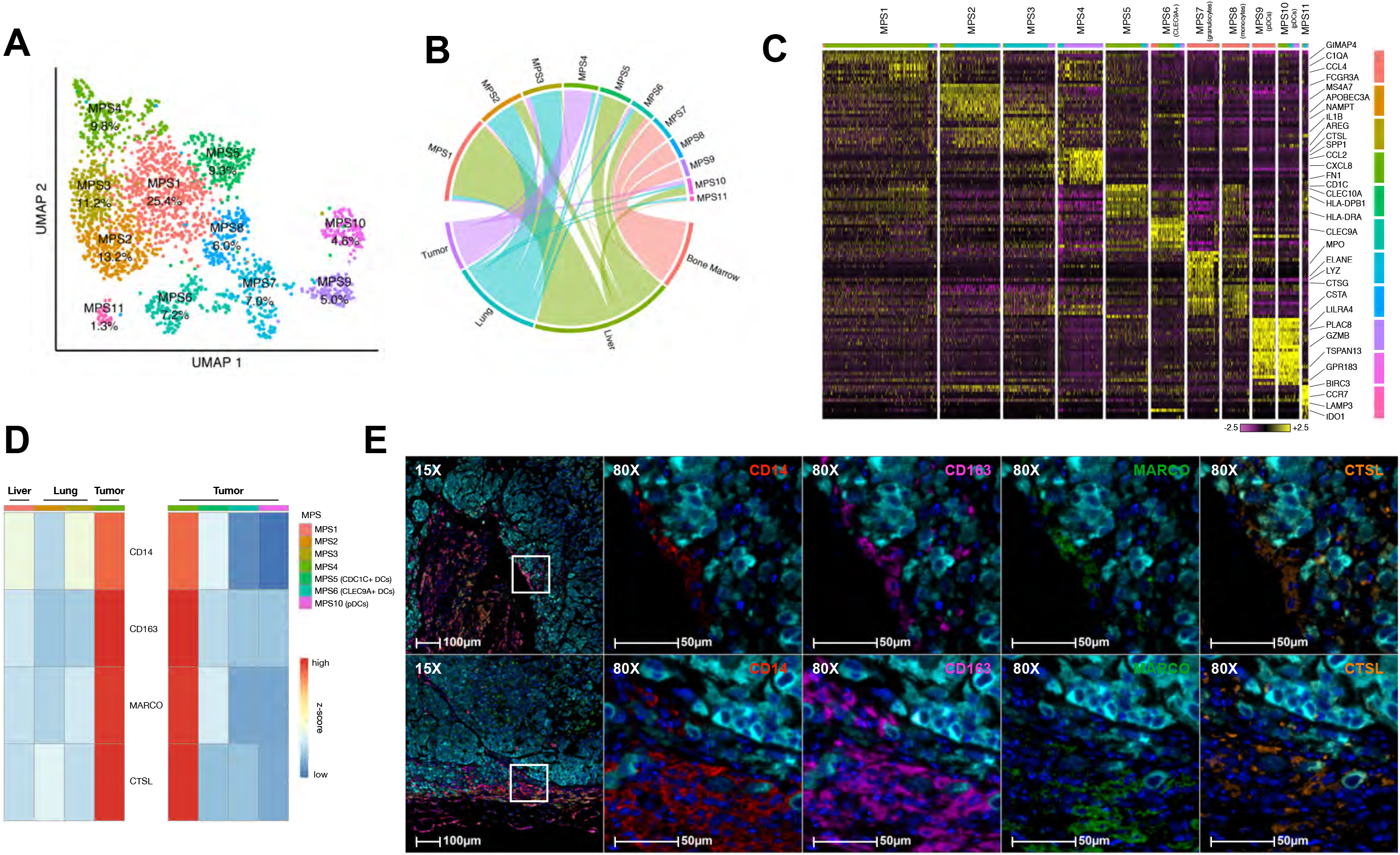
A distinct MPS cell cluster is present in the tumor microenvironment. (**A**) UMAP embedding of MPS cells (n = 2,152 cells) colored by unsupervised clustering. Identification of those clusters, based on expression of marker genes, is shown in Suppl. Fig 2B and Suppl. Table 2A. (**B**) Distribution of MPS clusters across tissues illustrated with circular plot. (**C**) Heatmap reporting scaled expression of the top 10 discriminative genes for each MPS cell cluster. The color scheme is based on z-score distribution from low (−2.5; purple) to high (+2.5; yellow). (**D**) Heatmap of expression of genes encoding target proteins selected for identification of MPS4 cells by mIHC. (**E**) Representative mIHC analysis of MPS4 cells infiltrating a Me275 melanoma in two individual MISTRG mice (melanoma, cyan; CD14, red; CD163 macrophages, magenta; MARCO, green, CTSL, orange).

### MPS4 cells are distinct from M2 polarized cells and myeloid-derived suppressor cells

Tumor-associated macrophages are frequently described in the context of the M1/M2 paradigm, which posits that M1 polarization, induced *in vitro* by IFNγ treatment, is associated with pro-inflammatory and anti-tumoral macrophage properties [30]. In contrast, IL-4-induced M2 polarization is associated with tissue-repair and tumor-supportive functions. *In vivo*, this framework has been extended to a proposed continuum of macrophage activation states, with M1 and M2 representing the extremes [31]. Besides M1/M2, comprehensive profiling has defined 49 transcriptional modules of human macrophage activation under 29 experimental *in vitro* conditions, associated with a multipolar spectrum of nine macrophage activation states [32]. We performed Gene Set Enrichment Analyses (GSEA) to determine the gene-level differences of these 49 transcriptional modules in MPS4 cells, compared to other non-DC tissue MPS cell clusters (**Figure 4A**). Only module 8 was significantly different in all tested comparisons, showing decreased expression in MPS4 cells (**Figure 4A, B**). Since module 8 was previously shown to be most highly expressed under canonical M1 stimulation with IFNγ [32], MPS4 cells have an apparent “anti-M1” transcriptional profile. Module 15 is associated with canonical M2 polarization by IL-4 [32] but only four genes from this module (*CD14, ALOX5AP, S100A8* and *S100A9*) were clearly overexpressed in MPS4 cells (**Figure 4A, B**). The predominantly liver-localized MPS1 cells had the most “M2-like” phenotype, with over-expression of modules 10 and 11 (**Figure 4A**). Finally, module 16 overexpression is associated with fatty acid response *in vitro* [32], and was driven by only three genes (*CTSD, GCHFR* and *LAMP1*) in MPS4 (**Figure 4A, B**). In addition, single-cell resolution heatmaps of all individual genes within the 49 transcriptional modules produced no obvious pattern, in contrast to the defining MPS4 ‘up’ and ‘down’ transcriptional signatures (**Suppl. Fig 3, Suppl. Table 2B**). Thus, MPS4 cells do not clearly correspond to any previously described transcriptional program. These results strongly support a reassessment of *in vivo* macrophage polarization paradigms [33], particularly in human cancer.

**Figure 4.**
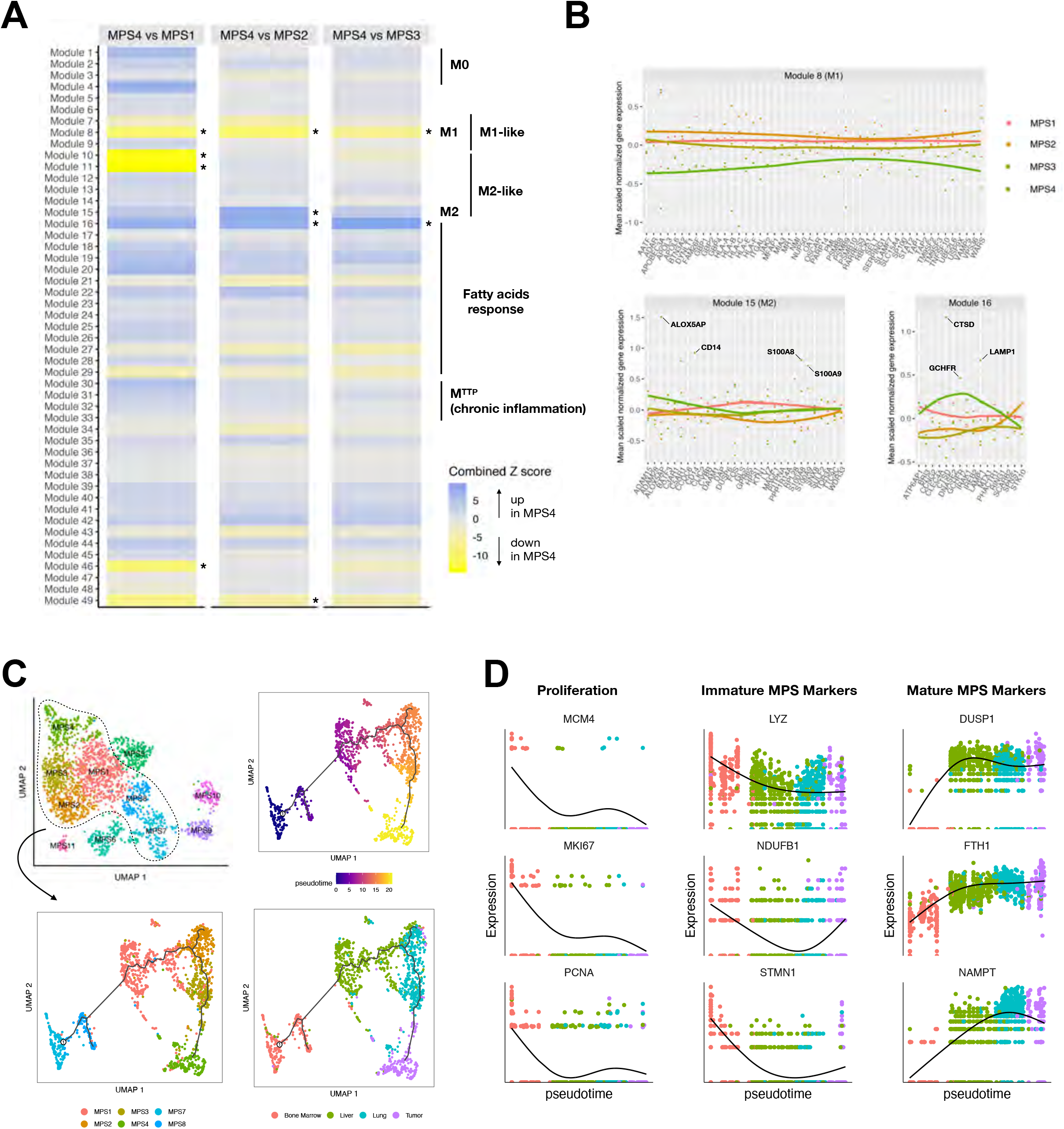
MPS4 cells do not correspond to canonical M2 macrophages nor MDSCs. (**A**) Heatmap of combined GSEA Z score using modules from Xue et al. [32] The combined Z-score reflects the enrichment due to differences in the continuous and discrete components of the single-cell MAST model between the three contrasts (MPS4 vs MPS1, 2 or 3). * indicate FDR 1%. (**B**) Mean scaled expression values of genes from module 8, 15 or 16. Genes are alphabetically ordered. Module 8 was declared significant in all the three contrasts (MPS4 vs MPS1, 2 or 3). (**C**) Pseudotime reconstruction of developmental trajectories in a two-dimensional independent component space using Monocle. Each dot represents a single cell, colored by pseudotime, MPS cluster or tissue of origin. (**D**) Expression patterns of key markers for proliferation, immaturity and maturity along pseudotime. Each dot represents a single cell, colored by tissues of origin. The x axis shows pseudotime and the y axis shows the relative expression of the markers. The black lines represent smooth expression curves.

Tumor-associated macrophages are also characterized within the framework of “myeloid-derived suppressor cells” (MDSCs) [34]; i.e., immature myeloid cells that harbor T cell inhibitory properties and accumulate in diverse conditions of non-resolving inflammation, such as chronic infection, autoimmunity or cancer. Moreover, MDSCs have been described as monocytic (M-MDSC) or polymorphonuclear (PMN-MDSC). Phenotypically, human MDSCs are poorly defined and cannot be fully discriminated from normal myeloid cells of corresponding lineages: M-MDSCs are reported as CD11b^+^ CD14^+^ HLA-DR^low^ CD15^-^ cells, and PMN-MDSCs as CD11b^+^ CD14^-^ CD15^+^ CD66b^+^ cells. Developmentally, MDSC populations have been described as myeloid progenitor and precursor cells that proliferate and accumulate but lack expression of terminal differentiation markers [35]. Based on high CD14 (**Figure 3D**) and low human leukocyte antigen (HLA) expression (described below), MPS4 cells could be labeled as M-MDSCs. However, pseudotime reconstruction of developmental trajectories of human MPS cells in MISTRG (**Figure 4C**) did not link MPS4 cells with this developmental definition of MDSCs. Indeed, MPS4 cells branched out from the MPS tree at an advanced pseudotime, suggesting that they derive from already mature cells. Furthermore, unlike BM progenitors, MPS4s showed no expression of proliferation markers (**Figure 4D**). Finally, we compared BM progenitors (MPS7 and 8) to differentiated cells in normal tissues (MPS1-3), with the objective of identifying transcriptional markers of MPS immaturity *vs*. maturity. Using these markers, MPS4 cells presented a clearly mature phenotype (**Figure 4D**), suggesting that they do not represent a population of immature or undifferentiated cells.

### Transcriptional network analysis reveals biological properties of MPS4 cells

To elucidate the biology of MPS4 cells beyond the M1/M2 and MDSC concepts, we performed GSEA comparing MPS4 (tumor) against MPS1 (liver) or MPS2 and MPS3 (lung), followed by network analysis (**Figure 5A, Suppl. Table 3**) [36]. Three gene sets stood out as upregulated in MPS4 cells; i.e., those associated with glucose metabolism, lysosome or cytokines/chemokines (**Figure 5A**). Most genes encoding enzymes involved in glycolysis and lactate production were overexpressed in tumor-associated MPS4 cells, compared to cells from other tissues (**Figure 5B,C**; **Suppl. Table 3D**). In contrast, gene sets encoding proteins involved in the tricarboxylic acid cycle and mitochondrial electron transport chain were downregulated (**Figure 5B, Suppl. Table 3D**). These data suggest an adaptation to the metabolically hostile and hypoxic TME, in which glucose is scarce and regeneration of the NAD^+^ pool is performed by reduction of pyruvate to lactate. Among immune regulators, the upregulation of several chemokines (CCL2, CXCL2, CXCL8) involved in the recruitment of myeloid cells to the TME suggests that a feed-forward mechanism shapes the immune composition of the TME. Other cytokines upregulated in MPS4 cells are known for their immunoregulatory functions (e.g., IL1RA encoded by *IL1RN*) or angiogenic role (VEGFA) in the TME (**Figure 5D**). Transcription factors, including HIF-1 pathway and AP-1 transcription factors, emerged as candidates bridging metabolism to expression of immunoregulatory cytokines, particularly the apparent switch in the *JUN*/*JUNB* ratio in MPS4 cells (**Figure 5E**). Finally, the overexpression of genes encoding lysosomal enzymes (such as cathepsins) suggests that MPS4 cells may degrade antigens, rather than process them for antigen presentation to T cells (**Figure 5F**).

**Figure 5.**
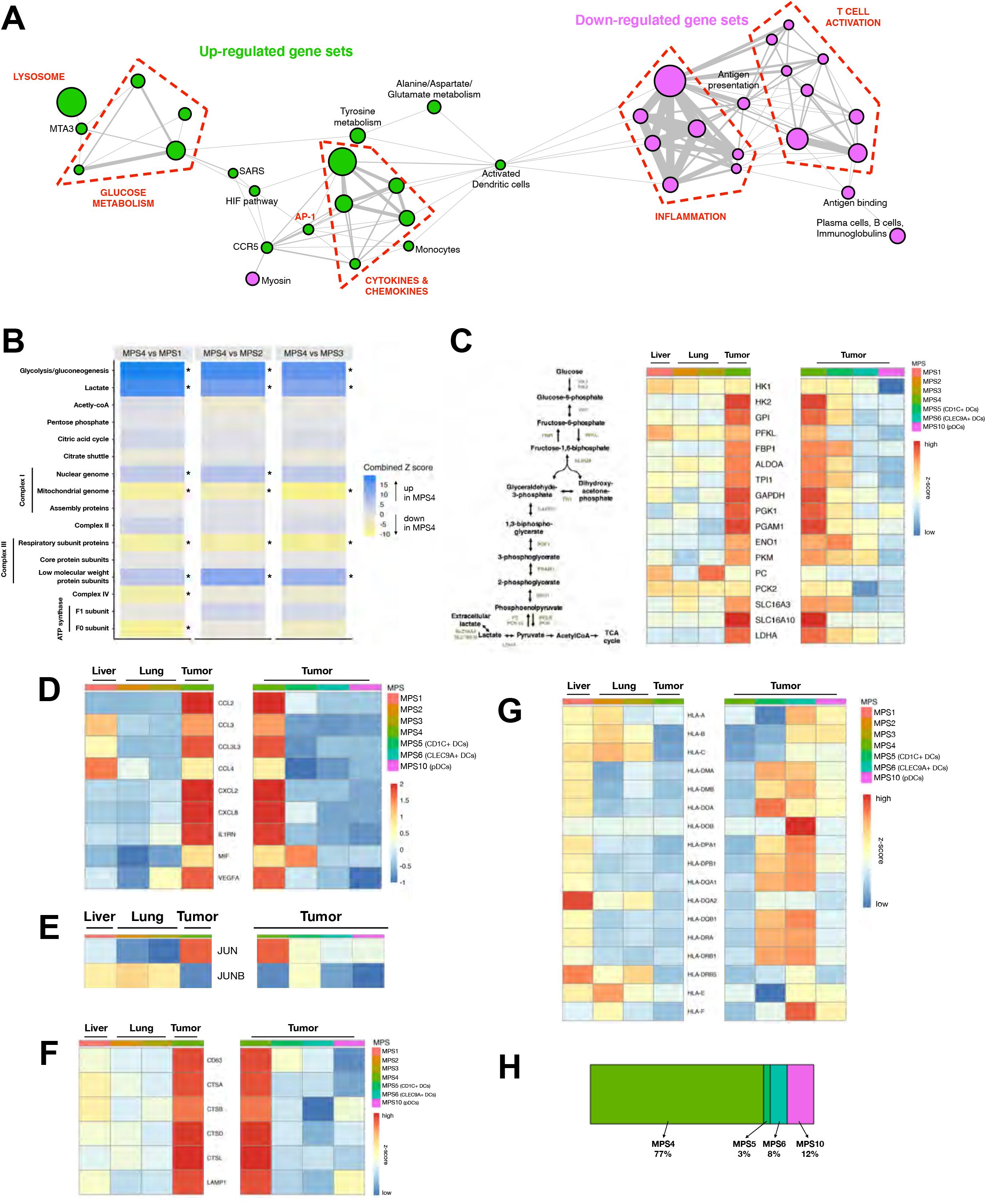
Transcriptional network analysis reveals biological properties of MPS4 cells. (**A**) Network representation of GSEA results. Each node represents a gene set differentially expressed in all three contrasts (MPS4 vs MPS1, MPS2 and MPS3), colored by either down (purple) or up (green) in MPS4. The size of the nodes is related to the number of genes in the gene set. Each edge indicates that two nodes share at least one gene. The width of the edges is related to the number of genes that two gene sets share. (**B**) Heatmap of combined GSEA Z score using modules of genes encoding enzymes involved in glycolysis and oxidative phosphorylation. The combined Z-score reflects the enrichment due to differences in the continuous and discrete components of the single-cell MAST model between the three contrasts (MPS4 vs MPS1, 2 or 3). * indicate FDR 1%. (**C**) Heatmap reporting the average expression across MPS1-4 clusters of genes encoding glycolytic enzymes. (**D**-**G**) Heatmap reporting the average expression across clusters of genes encoding chemokines/cytokines (**D**), the AP-1 family members JUN and JUNB (**E**), proteins associated with lysosome function (**F**), and HLA molecules (**G**) in macrophages (left) and dendritic cells in tumor (right). (**H**) Proportion of each MPS cluster among tumor-infiltrating cells.

Accordingly, antigen presentation- and other T cell activation-associated gene sets were downregulated in MPS4 cells (**Figure 5A**). In particular, the expression of all HLA molecules was low in these cells, compared to MPS from other tissues (**Figure 5G**). Multiple DC subsets were present in the TME (CD1C^+^ DCs, CLEC9A^+^ DCs and pDCs) and appeared to have more immunostimulatory properties than MPS4 cells, based on higher HLA and lower lysozyme enzyme expression (**Figure 5D-G**). However, these DC clusters were largely outnumbered by MPS4 cells in the TME (**Figure 5H**). These observations suggest a dominant role of MPS4 cells in shaping the composition and functional properties of the immune microenvironment in the tumor.

### The MPS4 transcriptional signature is expressed by cells infiltrating patient tumors

A humanized mouse model can be useful only if it accurately recapitulates human pathophysiology and is representative of diverse patient samples. To evaluate the broad applicability of the Me275 data, we generated a second humanized mouse melanoma model, using the A375 human melanoma cell line, and compared it to Me275 melanoma. Using mIHC, we found cells co-expressing the characteristic MPS4 markers (CD14, CD163, MARCO and CTSL) in both models (**Figure 6A**). scRNAseq analysis of tumor-infiltrating human CD45^+^ immune cells (**Figure 6B**) revealed that the Me275 TME is relatively enriched in MPS cells while the A375 contains more T cells (**Figure 6C**). However, all clusters perfectly overlaid in the UMAP across samples, suggesting similar transcriptomes. MPS cells (including pDCs) are distributed in 6 clusters in this analysis, labeled C8 to C13. We evaluated the expression levels of the MPS1-MPS11 gene sets, as defined in Suppl. Table 2A, by cells of each of these clusters. This analysis revealed that clusters C8 and C9 display a high expression of the MPS4 signature (**Figure 6D**). Together, C8 and C9 represent over 50% of all the MPS cells present in the TME of Me275 and A375 (**Figure 6E**).

**Figure 6.**
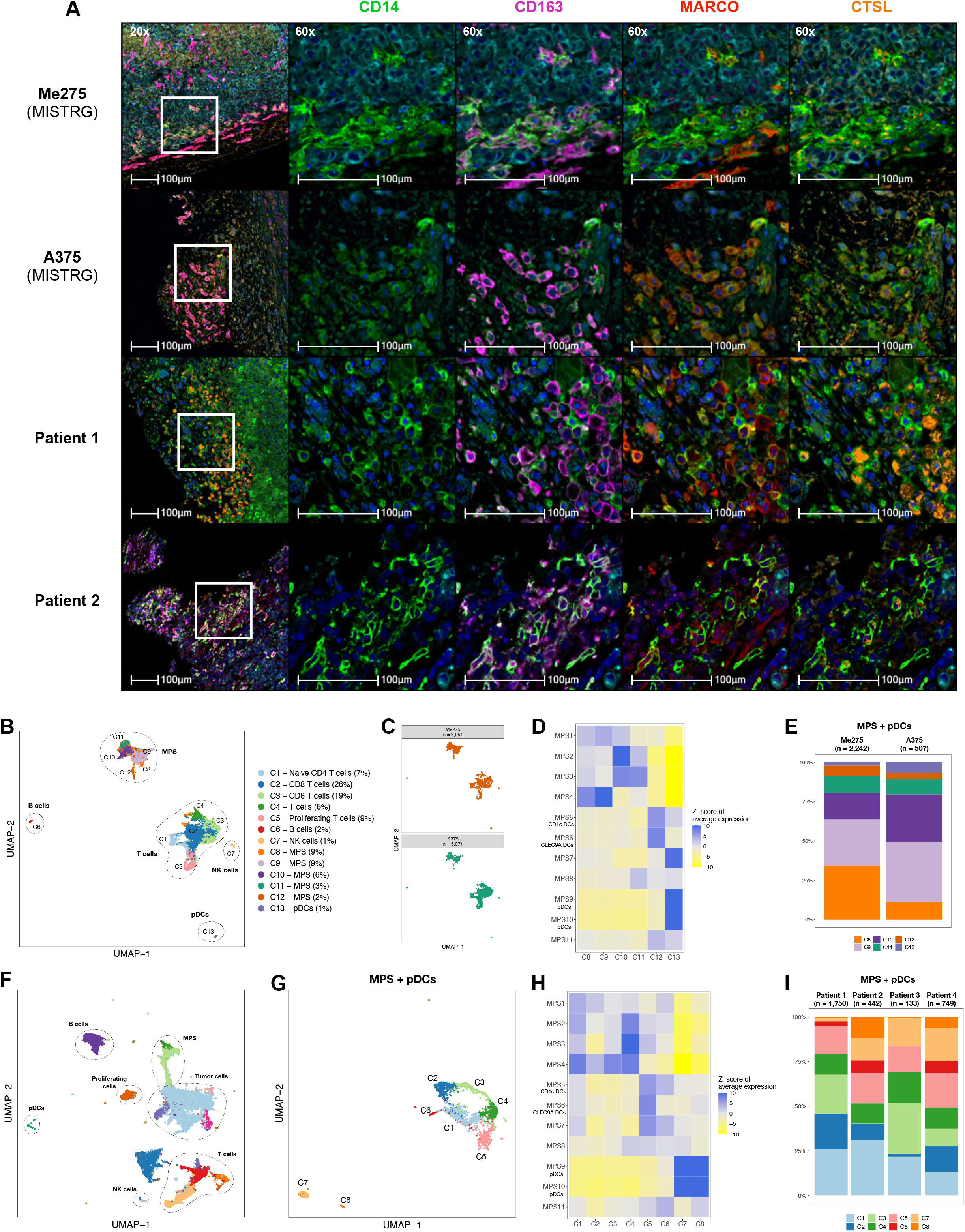
MPS4 cells are present in several humanized mouse and patient melanomas. (**A**) Representative images of the co-expression of MPS4 cell markers detected by multiplexed IHC in the TME of Me275 and A375 melanoma cell lines implanted in humanized MISTRG mice, and in the TME of two representative melanoma patients (melanoma, cyan; CD14, green; CD163, magenta; MARCO, red, CTSL, orange). See also **Suppl. Fig. 4** for additional melanoma patient samples. (**B, C**) UMAP representing the overlay (**B**) or distribution per tumor type (**C**) of human CD45^+^ cells isolated from the TME of Me275 and A375 tumors from humanized MISTRG mice and analyzed by scRNAseq. (**D, E**) Heatmap representing the expression of the MSP1-11 gene sets (as defined in Suppl. Table 2A; **D**), and relative distribution in each tumor (**E**) of clusters C8-C13 corresponding to MPS and pDC clusters delineated in **B**. (**F, G**) UMAP representing all sequenced cells (**F**) or MPS + pDCs (**G**) from single suspensions of tumors of four patient melanoma samples. (**H, I**) Heatmap of the expression of the MSP1-11 gene sets (**H**), and relative distribution in each patient (**I**) of clusters C1-C8, delineated in **G**.

We next characterized melanoma samples from seven patients with therapy-resistant melanoma undergoing surgical resections for metastatic disease. mIHC analysis revealed the presence of cells co-expressing the characteristic MPS4 markers in each of the samples (**Figure 6A, Suppl. Fig. 4**). We performed scRNAseq on cell suspensions prepared from four of the patient tumors (**Figure 6F**). To increase the resolution of our analysis, we re-clustered MPS cells (including pDCs) from the four patient samples (**Figure 6G**), and performed the same analysis as for the humanized mouse samples. The human clusters C1-C4 displayed the highest level of expression of the MPS4 transcriptional signature (**Figure 6H**), and these clusters contributed to 50-80% of MPS/pDC cells present in each individual patient sample. Together, these results show that cells transcriptionaly similar to MPS4 cells are present and abundant in the TME of human melanoma, across several humanized mouse models and patient samples.

A recent study generated a single-cell transcriptional atlas of tumor-infiltrating myeloid cells from over 200 patients and across 15 human cancer types, including melanoma [16]. This atlas revealed heterogeneous macrophage clusters, some shared across diverse cancer types. In melanoma, four tumor-infiltrating macrophage clusters were identified and named after their defining marker (Macro_VCAN, Macro_C1QC, Macro_ISG15 and Myeloid_MKI67); these clusters were distributed heterogeneously in different patient tumor samples but absent from circulating myeloid cells that were primarily Mono_CD14 and Mono_CD16 cells [16] (**Figure 7A, B**; **Suppl. Fig. 5A**). To determine whether any of the melanoma-infiltrating macrophage clusters corresponded to MPS4 cells, we examined whether they expressed the MPS4-defining gene set (**Suppl. Table 2A**). All four melanoma tumor-associated clusters showed a marked increase in MPS4-associated genes, compared to the other myeloid cell clusters (**Figure 7C**). Furthermore, these patient melanoma-infiltrating myeloid cells did not express any of the transcriptional signatures associated with MPS1-3 or MPS5-11 cells (**Suppl. Table 2A, Suppl. Fig. 5B**). These results demonstrate that the MPS4 gene set is expressed by heterogeneous, tumor-infiltrating macrophage cell clusters in melanoma patients. In contrast, the MPS4 gene set is not expressed by DCs or circulating monocytes (**Figure 7C, Suppl. Fig. 5B**).

**Figure 7.**
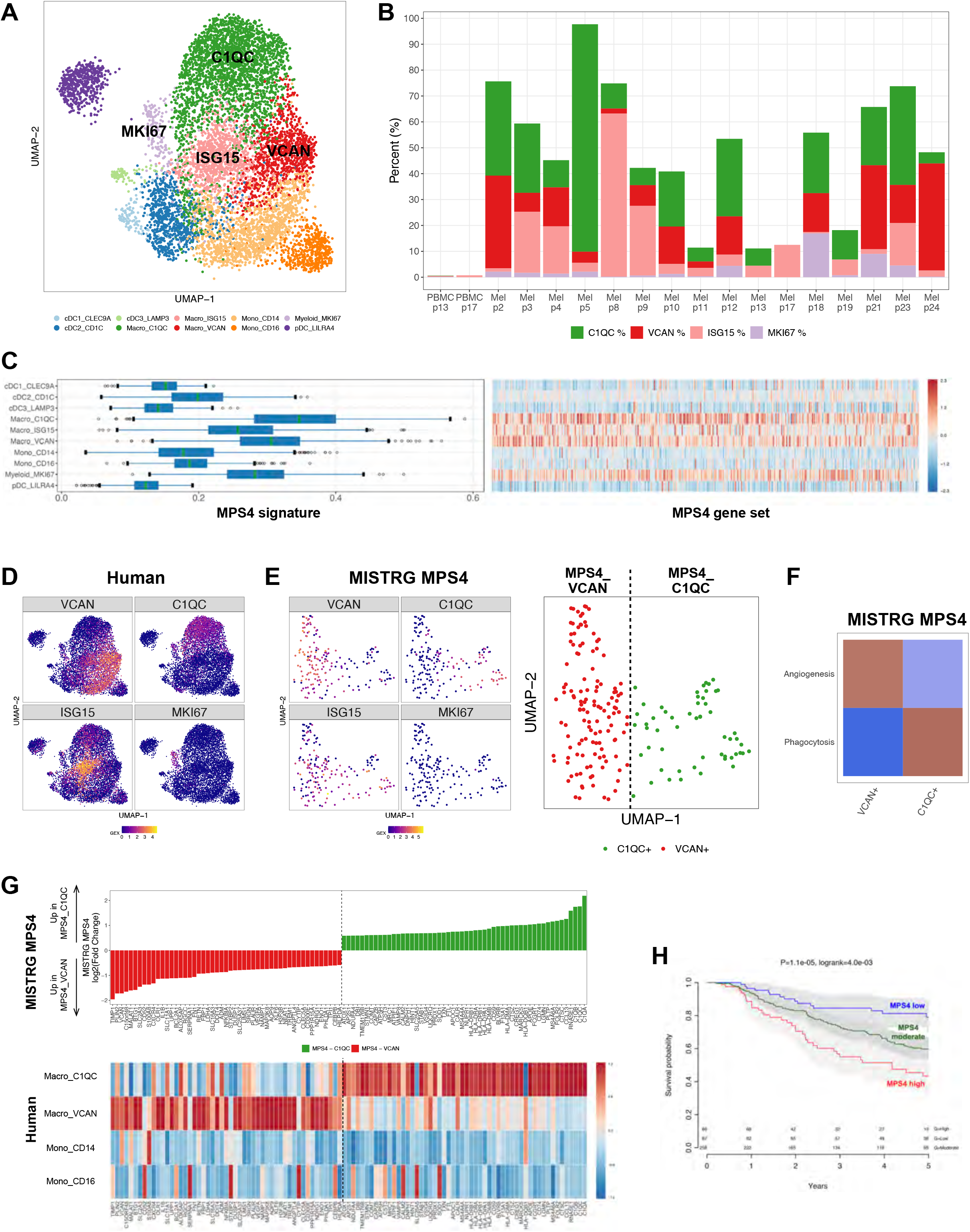
The MPS4 transcriptional signature corresponds to macrophages in a pan-cancer atlas of myeloid cell. (**A, B**) UMAP embedding of patient melanoma-infiltrating macrophages, dendritic cells and monocytes from publicly available scRNAseq data [52], previously analyzed and annotated by Cheng et al [16] (**A**). Distribution across patient samples (melanoma and PBMCs) of the Macro_C1QC, Macro_VCAN, Macro_ISG15 and Myeloid_MKI67 defined by Cheng et al [16] as melanoma-infiltrating macrophages (**B**). (**C**) Expression level of genes of the MPS4 cell transcriptional signature (from Suppl. Table 2) by cell clusters identified in **A** among cells infiltrating patient melanoma samples or control PBMCs. The MPS4 gene set is overexpressed in all four tumor-infiltrating macrophage clusters compared to all other cell clusters. (**D, E**) UMAP embedding of the expression of the defining markers (VCAN, C1QC, ISG15 and MKI67) among melanoma patient-infiltrating cells (**D**) and among MPS4 infiltrating the Me275 melanoma in humanized MISTRG mice (**E**, left). Manual re-clustering of MPS4 cells into MPS4_VCAN and MPS4_C1QC (**E**, right). (**F**) Heatmap showing the differential expression, between humanized mouse MPS4_VCAN and MPS4_C1QC, of the angiogenesis- and phagocytosis-associated gene sets (Suppl. Table 5A), identified as characteristic signatures of the corresponding human melanoma infiltrating Macro_VCAN and Macro_C1QC populations[16]. (**G**) Genes differentially expressed between humanized mouse MPS4_VCAN and MPS4_C1QC sub-clusters (top, Suppl. Table 5A), and heatmap representing the expression of these genes in patient Macro_VCAN and Macro_C1QC cells, in comparison to control CD14^+^ and CD16^+^ monocytes (bottom). (**H**) Kaplan-Meier plot stratified by high, intermediate or low overall expression of the MPS4 transcriptomic signature in bulk RNA-Seq of 473 skin cutaneous melanoma (SKCM) tumors from TCGA. P value: COX regression.

Because human melanoma-infiltrating macrophage subsets can be readily identified by the expression of their defining marker [16] (**Figure 7D**), we wanted to determine whether we could identify a similar heterogeneity among MPS4 cells from humanized mice. *ISG15* was expressed at similar levels among all MPS4 cells, and *MKI67* was barely detectable (**Figure 7E**). However, we could distinguish mutually exclusive expression of *VCAN* and *C1QC* in two subsets of cells. Therefore, we manually divided MPS4 cells into MPS4_C1QC and MPS4_VCAN sub-clusters (**Figure 7E**). In patient samples, human Macro_VCAN and Macro_C1QC have been associated with the expression of gene sets related to angiogenesis and phagocytosis, respectively [16] (**Suppl. Table 4A**). Recapitulating these results from patient samples, MPS4_VCAN and MPS4_C1QC from humanized mice were also associated with expression of the corresponding gene sets (**Figure 7F**). We next performed the opposite analysis, identifying a list of genes differentially expressed between humanized mouse MPS4_VCAN and MPS4_C1QC (**Figure 7G** top, **Suppl. Table 4B**) and determined the expression of these gene sets among corresponding cell clusters found infiltrating patient melanomas. We found a good concordance between human and humanized mouse VCAN^+^ and C1QC^+^ subsets (**Figure 7G**).

After establishing the MPS4 transcriptional signature in cells infiltrating patient melanoma samples, we assessed whether it was associated with disease outcomes. For this, we examined MPS4-associated up and down gene expression (**Suppl. Table 2B**) in bulk RNA-seq data from 473 melanoma patient samples in the Cancer Genome Atlas (TCGA), using the Overall Expression algorithm [37]. This analysis associated high level expression of the MPS4 transcriptional profile with significantly shorter overall survival of melanoma patients (**Figure 7H**).

Finally, a broad SPP1-expressing macrophage population is abundant across diverse tumor types, beyond melanoma, and these cells highly express the same angiogenesis transcriptional signature as melanoma Macro_VCAN^+^ cells [16]. In fact, although melanoma-infiltrating macrophages were not previously labeled as Macro_SPP1 cells [16], we found SPP1 highly expressed in both VCAN^+^ and C1QC^+^ cells infiltrating melanomas in human samples and humanized mice (**Suppl. Fig 6A**). Thus, we determined whether the MPS4 gene set (**Suppl. Table 2**) was also expressed in SPP1^+^ macrophages infiltrating other human cancer types. We focused our analysis on colorectal cancer (CRC), thyroid carcinoma (THCA) and uterine corpus endometrial carcinoma (UCEC), for which data is available for both tumor and adjacent non-tumor tissue [16]. We confirmed that the Macro_SPP1 cluster was enriched in all three tumor types, compared to control tissues (**Suppl. Fig 6B**), and most abundantly expressed the highest level of the MPS4 transcriptional signature across all three cancer types (**Suppl Fig 6C**). Taken together, these observations demonstrate that MPS4 cells, identified in a humanized mouse model of metastatic melanoma, share the transcriptional characteristics of macrophage populations that broadly infiltrate human cancers.

## Discussion

The exciting but differential successes of recently advanced cancer immunotherapies demonstrate the importance of better understanding the functional interactions between multiple cell types in TMEs. In particular, cancer-immune cell interactions have definitive impacts on therapeutic responses and long-term patient outcomes. Transformative technologies, such as scRNAseq, are providing comprehensive surveys of the heterogenous composition of immune TMEs in human cancers [38], including the heterogeneity of human myeloid cells [39]. This newly-generated descriptive knowledge provides novel perspectives on how we could design innovative therapeutic strategies, which could target immune cells of the TME that impact disease progression. However, experimental models are needed to faithfully recapitulate the functional properties of immune cells in human cancer, to enable experimental perturbation of the immune TME, testing of new hypotheses developed from scRNAseq results, and ultimately a better understanding of the fundamental mechanisms underlying immune-cancer cells interactions. These models are also needed to evaluate new candidate therapies under clinically-relevant conditions.

Here, we demonstrate that the humanized MISTRG immuno-CDX model of Me275 melanoma recapitulates the tumor-supportive function of macrophages in human cancer, particularly their role in facilitating metastatic tumor spread. We report an in-depth scRNAseq characterization of tumor-infiltrating MPS cells, revealing a unique transcriptional signature of tumor-infiltrating “MPS4” cells versus MPS cells in other tissues, which allows us to infer the functional properties of these cells from their transcriptome. While the M1/M2 polarization has been widely used to describe macrophage function in cancer, this paradigm only partially represents the transcriptome of MPS4 cells. Tumor-infiltrating myeloid cells are also described as MDSCs, and this concept broadly describes myeloid cells associated with non-resolving inflammation in a number of pathologies, but unequivocal MDSC markers are not clearly identified. The MPS4 cell transcriptome is only partially concordant with the described properties of MDSCs; the absent expression of HLA molecules and gene sets associated with T cell activation is compatible with the immunosuppressive properties of MDSCs, but the reconstruction of MPS4 developmental trajectories contrasts with the immature nature of MDSCs.

Our studies show that the MPS4 transcriptional signature is highly similar to those of macrophage populations identified in human melanoma and other cancer types. In human melanoma, the VCAN^+^ and C1QC^+^ macrophage populations highly express the MPS4 gene set. In addition, a re-analysis of humanized mouse MPS4 cells, based on macrophage heterogeneity identified in patient melanomas, reveals that similar subsets are found in humanized mice and recapitulate the differential expression of pro-angiogenesis vs. phagocytosis transcriptional signatures. Among macrophages infiltrating other human tumors, the SPP1^+^ macrophage subset notably expresses the MPS4 gene set at high levels. Therefore, knowledge gained from the humanized mouse melanoma model might be applicable to other cancer types, although additional validations will be needed.

The similarities between human and humanized mouse melanoma-infiltrating macrophages demonstrate the complementarity between these two experimental systems—patient sample analyses provide high-dimensional descriptive characterizations of immune TMEs, while validated humanized mouse models enable experimental perturbations. Together, these combined approaches have the potential to enhance our fundamental understanding of tumor ecosystems, and enable the development and pre-clinical testing of rational candidate therapies under clinically-relevant conditions.

However, some limitations remain. In particular, antigen-specific adaptive immune responses are weak and/or delayed in all available humanized mouse models [21, 40]. Consequently, evaluation of checkpoint inhibition therapies in humanized mice is complicated and frequently produces highly variable results with different human donors [41]. Novel humanized mouse models, with improved adaptive immunity, are showing some promise [42] and approaches are being developed to employ the adoptive transfer of T cells with pre-determined antigen specificity (e.g. transgenic chimeric antigen receptor or T cell receptor T cells). Our model provides unique opportunities for *in vivo* studies of multiple aspects of macrophage biology in human cancers, based on the most recent knowledge gained from single cell transcriptomics. Together, these technological advances will hopefully lead to the development of new, life-saving therapies for cancer patients.

## Material and methods

### Mice

MISTRG mice (<u>M</u>-CSF^h/h^ <u>I</u>L-3/GM-CSF^h/h^ <u>S</u>IRPα^h/m^ <u>T</u>PO^h/h^ <u>R</u>AG2^-/-^ IL-2Rγ^-/-^) were previously reported [22, 43]. In these mice, several cytokine-encoding genes (*Csf1, Il3/Csf2* and *Thpo*) are humanized by knockin replacement of the mouse allele by its human ortholog from ATG to stop codon, in a *Rag2 Il2rg* double knockout 129xBALB/c (N2) background. The *Sirpa* gene is humanized by replacement of the sequence encoding the extracellular domain of SIRPα. MISTRG mice were obtained from Yale University and are used under Material Transfer Agreements with Yale University and Regeneron Pharmaceuticals. ISTRG mice (<u>I</u>L-3/GM-CSF^h/h^ <u>S</u>IRPα^h/m^ <u>T</u>PO^h/h^ <u>R</u>AG2^-/-^ IL-2Rγ^-/-^) retain the wildtype M-CSF^m/m^ allele, but are otherwise identical to MISTRG. To generate heterozygous MIS^h/m^TRG mice for human cell transplantation, we intercross homozygous MIS^h/h^TRG males and MIS^m/m^TRG females that we maintain as distinct colonies, thus avoiding any genotyping requirement. We are using a similar strategy to generate IS^h/m^TRG recipient mice. After rederivation by embryo transfer, all mice were housed in an enhanced barrier (with restricted access and enhanced personal protective equipment requirements) under specific pathogen free conditions, with continuous prophylactic enrofloxacin treatment (Baytril, 0.27 mg/ml in drinking water). All animal experiments were approved by Fred Hutchinson Cancer Research Center (Fred Hutch)’s Institutional Animal Care and Use Committee (protocol # 50941).

### Human CD34^+^ cell isolation and transplantation

De-identified human fetal liver tissues, obtained with informed consent from the donors, were procured by Advanced Bioscience Resources, Inc. and their use was determined as non-human subject research by Fred Hutch’s Institutional Review Board (6007-827). Fetal livers were cut in small fragments, treated for 45 min at 37°C with collagenase D (Roche, 100 ng/ml), and a cell suspension was prepared. Hematopoietic cells were enriched by density gradient centrifugation (Lymphocyte Separation Medium, MP Biomedicals) followed by positive immunomagnetic selection with anti-human CD34 microbeads (Miltenyi Biotec). Purity (>90% CD34^+^ cells) was confirmed by flow cytometry and cells were frozen at -80°C in FBS containing 10% DMSO. Newborn mice (day 1-3) were sublethally irradiated (80 cGy gamma rays in a Cesium-137 irradiator) and ∼20,000 CD34^+^ cells in 20 μl of PBS were injected into the liver with a 22-gauge needle (Hamilton Company), as previously described [44, 45]. Engraftment levels were measured as the percentage of human CD45^+^ cells among total (mouse and human combined) CD45^+^ cells in the blood.

### Flow cytometry analysis

To evaluate the engraftment levels and multilineage engraftment of human hematopoietic cells in mice, blood was obtained by retro-orbital collection and red blood cells were eliminated by ammonium-chloride-potassium (ACK) lysis. White blood cells were analyzed by flow cytometry, following standard procedures. The following antibody clones were used (all purchased from Biolegend):

Anti-human antibodies: CD3-AF700 (HIT3a), CD19-PE-Cy7 (HIB19), CD33-APC (WM53), CD34-PE (581), CD45-Pacific Blue (HI30), NKp46-PE (9E2).

Anti-mouse antibodies: CD45-BV605 (30-F11), Ter119-PerCP (TER-119).

Human hematopoietic cells were gated based on expression of human CD45 and exclusion of mouse CD45 and Ter119 staining. Dead cells were excluded by staining with 7-Aminoactinomycin D (7-AAD)

### Tumor cell implantation and monitoring

The human melanoma cell line Me275 was obtained from Dr. P. Romero and used under a Material Transfer Agreement with the University of Lausanne (Switzerland), and the A375 cell line was obtained from the American Type Culture Collection (ATCC). Cells were grown to ∼80% confluency in DMEM supplemented with 10% fetal bovine serum. Approximately 7 million cells (Me275) or 1 million cells (A375) per mouse were injected subcutaneously under anesthesia in the flank of humanized the mouse. The size of the tumors was measured weekly for 7 weeks with a caliper, and the tumor volume was calculated using the following formula: volume = 0.5 × length^2^ × width. At the endpoint, metastatic nodules in the liver and spleen were enumerated. Tissues were processed for human CD45^+^ cell isolation and scRNA sequencing, or fixed in formalin (neutral buffered, 10%) for subsequent histology analysis.

### Patient samples

All clinical sample collections were conducted according to the Declaration of Helsinki principles. The protocol (FHCR 1765) to obtain biological samples was approved by the Fred Hutch’s Institutional Review Board. All patients signed informed consent prior to donating tumor tissue. Surgeries were clinically indicated and only leftover tumor material was donated to the study. The surgically removed tumor samples were sectioned by a pathologist in the operating room. A portion of the tumor was fixed in formalin for histology analysis; and another one was dissociated, cryopreserved and used for scRNAseq analysis.

### Human CD45^+^ cell isolation from MISTRG and scRNA sequencing

Tumors were dissected 3 weeks after implantation. BM, liver and lung were harvested from naïve mice. Tissues were pooled from 5 mice, previously repopulated with human CD34^+^ HSPCs from two individual donors.

Tumors were digested with Miltenyi’s Tumor Dissociation Kit, Human (Miltenyi Biotec) and human CD45^+^ cells sorted by flow cytometry on a FACS Aria II instrument.

BM cells were treated with ACK to lyse red blood cells. Livers and lungs were treated with collagenase D (for lung digestion, 1 mg/ml, Roche) or collagenase Type 4 (for liver digestion, 1 mg/ml, Worthington), supplemented with DNase (10 μg/ml) for 30 minutes. White blood cells were separated on a Percoll gradient (Cytiva), and human CD45^+^ cells were selected by magnetic cell sorting (Miltenyi Biotec). scRNAseq libraries were prepared with the 10x Genomics’ Chromium Single Cell 3’ Library & Gel Bead Kit v2, following the manufacturer’s protocol, and sequenced on the Illumina HiSeq2500 instrument.

### scRNAseq data analysis – MISTRG Me275 vs. tissues

#### Transcriptome alignment, barcode assignment and UMI counting

The Cell Ranger Single-Cell Software Suite (version 2.0.1) was used to perform demultiplexing, barcode processing and single-cell 3’ (5’) gene counting of samples (http://10xgenomics.com/). First, raw base BCL files were demultiplexed using the cellranger mkfastq pipeline into sample-specific FASTQ files. Second, these FASTQ files were processed with the cellranger count pipeline. Each sample was analyzed separately. The cellranger count pipeline uses the STAR alignment software to align cDNA reads to the pre-built GRCh38 human reference genome given by 10x Genomics. Aligned reads were then filtered for valid cell barcodes and unique molecular identifiers (UMIs). Cell barcodes with 1-Hamming-distance from a list of known barcodes were considered. UMIs with sequencing quality score > 10% and not homopolymers were retained as valid UMIs. A UMI with 1-Hamming-distance from another UMI with more reads, for a same gene and a same cell was corrected to this UMI with more reads. MISTRG samples (bone marrow, liver, lung and tumor) were aggregated together using the cellranger aggr pipeline resulting in one gene-barcode count matrix ready for analysis. Human melanoma samples (from two different patients) were also aggregated together using the same pipeline as above. A correction for sequencing depth was performed during the aggregation [26].

#### Data normalization

Only genes with at least one UMI count detected in at least one cell are used [26]. A library-size normalization was performed for each cell. UMI counts were scaled by the total expression in each cell and multiplied by 10,000. The data were then log-transformed before analysis as an input to Seurat [46].

#### All tissues analysis

Following sequence alignment and filtering, a total of 6,399 cells were analyzed (after removing cells that have unique gene counts less than 200, percentage of mitochondrial genes more than 20% or more than 40,000 UMIs) (**Suppl. Fig 1A**). The normalized gene-barcode matrix was used to run principal component analysis (PCA), clustering and uniform manifold approximation and projection (UMAP) analyses.

The top 1,790 variable genes were used as inputs to compute PCA. The first top 15 principal components (PCs) were used for UMAP visualization and clustering. UMAP with a min.distance=0.75 was performed (**Figure 2A** and **Suppl. Fig 1B**). Cell classification and clustering were performed using a graph-based clustering method implemented in Seurat (FindClusters R function - share nearest neighbor (SNN) modularity optimization based clustering algorithm). Briefly, it calculates k-nearest neighbors and constructs the SNN graph. Then, it optimizes the modularity function to determine the best clusters. Eighteen distinct clusters of cells were identified using the top 15 PCs with a neighborhood size of 30 and resolution of 0.6. Specific enriched markers for each cluster were identified using the FindAllMarkers R function (with test.use=‘MAST’ and latent.vars=‘nUMI’) (**Suppl. Table 1A** and **Suppl. Fig 1B**). Clustering and annotation of mononuclear phagocytic system (MPS) cells were then refined by selecting clusters composed of MPS (**Suppl. Fig 2A**). The top 849 most variable genes selected by Seurat were kept to compute PCA. As above, cell classification and clustering were performed using a graph-based clustering method implemented in Seurat. Eleven distinct clusters of cells were identified using the top 20 PCs with a neighborhood size of 25 and resolution of 0.75. FindAllMarkers R function was used to detect enriched markers for each cluster (**Suppl. Table 2A** and **Suppl. Fig 2B**).

#### Tumor sample analysis

This sample was analyzed following the same pipeline as above. A total of 477 cells were analyzed (cells with at least 200 unique genes, percentage of mitochondrial genes less than 15% or less than 30,000 UMIs). The normalized gene-barcode matrix was used to run principal component analysis (PCA), clustering and uniform manifold approximation and projection (UMAP) analyses using Seurat.

The top 3,511 variable genes (with log-mean expression values greater than 0.0125 and dispersion (variance/mean) greater than 0.5) were used as inputs to compute PCA. The first top 10 principal components (PCs) were used for UMAP visualization. UMAP with a min.distance=0.5 was performed. Ten distinct clusters of cells were identified using the top 10 PCs with a neighborhood size of 10 and resolution of 0.6. Specific enriched markers for each cluster were identified using the FindAllMarkers R function (with test.use=‘MAST’ and latent.vars=‘nUMI’) (**Suppl. Table 1B** and **Suppl. Fig 1C**).

#### Differential expression analysis

Differential expression analysis (DEGs and GSEA) between clusters or tissues were performed using the R package MAST [36]. Normalized gene-barcode matrices were always used as inputs. A logistic regression model is used to test differential expression rate between groups, while a Gaussian generalized linear model (GLM) describes expression conditionally on non-zero expression estimates. The model was also corrected for the cellular detection rate (CDR). Genes were declared significantly differentially expressed with FDR 1% and absolute log2-fold-change > log2(1.5). Gene sets were declared significant with FDR 1%, absolute continuous Z score > log2(2.5) and absolute discrete Z score > log2(2.5). Blood transcriptome modules (BTM) [47], KEGG and BioCarta pathways were used as gene sets. Those gene sets (KEGG and BioCarta) were downloaded from MSigDB database [48].

#### Technical effects

It is known that technical parameters such as the library size (total mapped UMIs) or the total number of genes detected can vary across cells. We used the specific ScaleData R function implemented in Seurat to remove technical effect due to the library size (total mapped UMIs) as well as the percentage mitochondrial gene content. The corrected gene-barcode matrix was used as input for PCA, UMAP and clustering analyses. The number of mapped UMIs per cell (used as proxy for CDR) was taken into account in the MAST analyses to get differentially expressed genes and gene sets.

#### Developmental pseudotime analysis

We used the Monocle3 package in R to order cells into a developmental pseudotime [49-51]. Following the Monocle vignette, we used UMI count data as input and selected genes declared as differentially expressed between clusters for ordering of the cells. The default settings were used for all other parameters.

#### Gene set variation analysis

Estimation of GSEA scores in Figure 7F were performed with the GSVA R package.

### scRNAseq data analysis – MISTRG Me275 vs. A375

#### Transcriptome alignment, barcode assignment and UMI counting

The Cell Ranger Single-Cell Software Suite (version 2.0.1) was used to perform demultiplexing, barcode processing and single-cell 3’ (5’) gene counting of samples as described above.

#### Statistical analyses

A relaxed QC strategy was used, and we chose to only exclude cells with large mitochondrial proportions as proxy for cell damage. A stringent cutoff (5 Median Absolute Deviation (MAD)) for percent of mitochondrial genes was used, resulting in a data set of 9,002 cells for downstream analyses (1,484 cells were filtered out). Only genes with at least one UMI count detected in at least three cells were kept (23,351 genes).

Data integration and standard analyses were performed using the Seurat R package (Bulter et al., 2019). Gene expression data integration was performed using the tumor as factor, resulting in two datasets. To integrate the gene expression values, we separately normalized each of our partitions using the *SCTransform* R function (Hafemeister et al., 2019) with *vars*.*to*.*regress = “percent*.*mito”*. Data integration was then performed using the two R functions *FindIntegrationAnchors* and *IntegrateData* (with *dims = 1:50*). Principal component analysis (PCA) on the integrated dataset was performed using the *RunPCA* R function. The top 50 principal components (PCs) were then used for UMAP visualization and clustering. UMAP with this parameter (*min*.*dist =* .*01*) was performed using the *RunUMAP* R function. Cell clustering was performed using a graph-based clustering method implemented in Seurat (*FindNeighbors* and*FindClusters* R function) with different resolutions. Cluster annotation was performed using the R package MAST (Finak et al., 2015) implemented within the *FindMarkers* R function from the Seurat R package. We merged clusters that do not exhibit clear evidence of separation based on differential expressed markers and visualization.

### scRNAseq data analysis – Melanoma patient samples

#### Transcriptome alignment, barcode assignment and UMI counting

The Cell Ranger Single-Cell Software Suite was used to perform demultiplexing, barcode processing and single-cell 3’ (5’) gene counting of samples as described above.

#### Statistical analyses

The same statistical pipeline as above was investigated. Again, a relaxed QC strategy was used. A cutoff (5 MAD) for percent of mitochondrial genes was used, resulting in a data set of 34,945 cells for downstream analyses (4,473 cells were filtered out). Only genes with at least one UMI count detected in at least three cells were kept (23,021 genes).

Data integration and standard analyses were performed using the Seurat R package (Bulter et al., 2019). Gene expression data integration was performed using the patient id as factor, resulting in four datasets to align. The same data integration approach, as well as visualization and clustering, as described above was used. Cluster annotation was also performed using the R package MAST (Finak et al., 2015) implemented within the *FindMarkers* R function from the Seurat R package. We merged clusters that do not exhibit clear evidence of separation based on differential expressed markers and visualization.

### Comparison to human atlas

Humanized mouse MPS cell transcriptional signatures were compared to scRNAseq data from a pan-cancer transcriptional atlas of tumor-infiltrating myeloid cells [16]. The analyses were performed on the http://panmyeloid.cancer-pku.cn/ portal.

### TCGA

We looked at the gene signature specific of MPS4 in the bulk RNA-seq of 473 skin cutaneous melanoma (SKCM) tumors from TCGA. Using the R code and data provided by Jerby-Arnon et al. [37] (https://github.com/livnatje/ImmuneResistance), we computed the overall expression of the lists of up (40 genes) and down (32 genes) differentially expressed genes between MPS4 and MPS1, 2 and 3 (**Suppl. Table 2B**); and predicted the overall survival in TCGA melanoma patients using this signature. Kaplan-Meier (KM) curves were stratified by high (top 20%), low (bottom 20%), or intermediate (remainder) overall expression.

### mIHC

Formalin-fixed paraffin-embedded tissues were sectioned (4µM) and stained on a Leica BOND Rx autostainer using the Akoya Opal Multiplex IHC assay (Akoya Biosciences, Menlo Park, CA) with the following changes: Additional high stringency washes were performed after the secondary antibody and Opal fluor applications using high-salt TBST (0.05M Tris, 0.3M NaCl, and 0.1% Tween-20, pH 7.2-7.6). TCT was used as the blocking buffer (0.05M Tris, 0.15M NaCl, 0.25% Casein, 0.1% Tween 20, pH 7.6 +/-0.1). All primary antibodies were incubated for 1 hour at room temperature.

The following staining panels were used:

| Position | Antibody | Clone / Host | Company / Item | OPAL Fluor |
| --- | --- | --- | --- | --- |
| 1 | CD3 | SP7 / Rabbit | Thermo Fisher / RM-9107 | 620 |
| 2 | Melanoma // Sox10 cocktail | M2-7C10; M2-9E3, T311, HMB45 / Mouse // Polyclonal / Rabbit | Novus / NBP2-34337 // CellMarque / 383A-74 | 570 |
| 3 | CD163 | EP324 / Rabbit | Bio SB / BSB3276 | 520 |
| 4 | CLEC9A | EPR22324 / Rabbit | Abcam / ab223188 | 690 |
| Secondary | Opal Polymer HRP Ms+Rb |  | Akoya Biosciences / ARH1001EA |  |
| 1 | Melanoma // Sox10 cocktail | M2-7C10; M2-9E3, T311, HMB45 / Mouse // Polyclonal / Rabbit | Novus / NBP2-34337 // CellMarque / 383A-74 | 650 |
| 2 | Cathepsin-L | 33/2 / Mouse | AbCam / ab197278 | 570 |
| 3 | CD14 | EPR3653 / Rabbit | Cell Marque / 114R-14 | 540 |
| 4 | CD163 | EP324 / Rabbit | Bio SB / BSB3276 | 690 |
| 5 | MARCO | Polyclonal / Rabbit | Sigma / HPA063793 | 620 |
| Secondary | Opal Polymer HRP Ms+Rb |  | Akoya Biosciences / ARH1001EA |  |

Slides were mounted with ProLong Gold and cured for 24 hours at room temperature in the dark before image acquisition at 20x magnification on the Akoya Vectra 3.0 Automated or Polaris Automated Imaging System. Images were spectrally unmixed or autofluorescence-subtracted using Akoya Phenoptics inForm software.

Images were analyzed using the HALO software.

### Statistical analyses

Statistical analyses were performed in GraphPad Prism, using the non-parametric Mann-Whitney test.

## Data availability

Single-cell RNA sequencing data submitted to National Center for Biotechnology Information Gene Expression Omnibus (NCBI GEO), accession GSE173625.

R code to reproduce the Seurat analyses is available on GitHub https://github.com/ValentinVoillet/MISTRG

## Acknowledgments

We thank lab and Department members for valuable discussions; Deborah Banker for manuscript editing and Silvia Christian for administrative support. We are grateful to Advanced Bioscience Resources, Inc. for providing fetal tissues; and Fred Hutch’s Comparative Medicine for outstanding mouse husbandry. We acknowledge Yale University, the University of Zürich and Regeneron Pharmaceuticals where MISTRG mice were generated thanks to the financial support of the Bill and Melinda Gates Foundation. This work was funded by the National Cancer Institute of the National Institutes of Health (R01 CA234720 to AR; T32 CA080416, to TRB), Safeway Albertsons, The Hartwell Foundation, Fred Hutch’s Immunotherapy Integrated Research Center and the Bezos family (to AR).

This research was also supported by Shared Resources (Comparative Medicine, Flow Cytometry, Genomics and Experimental Histopathology) of the Fred Hutch/University of Washington Cancer Consortium (P30 CA015704).

## Authorship Contributions

VV performed scRNAseq data analysis. TRB performed and analyzed in vivo experiments. KMM performed in vivo experiments and generated scRNAseq librariw. KGP obtained patient samples analyzed mIHC assays. KSS, SW, JSC and RHP developed mIHC assays. DSH coordinated patient sample acquisition and generated scRNAseq libraries. WJV and JHB participated to the design and execution of scRNAseq experiments. TSK and DRB performed patient surgeries and procured human melanoma samples. AGC supervised patient sample acquisition and analysis. RG supervised scRNAseq data analysis. VV and AR prepared the manuscript, and all authors provided feedback. AR conceived the study; designed, executed and interpreted experiments; and supervised research.

---

**Supplementary Figure 1.**
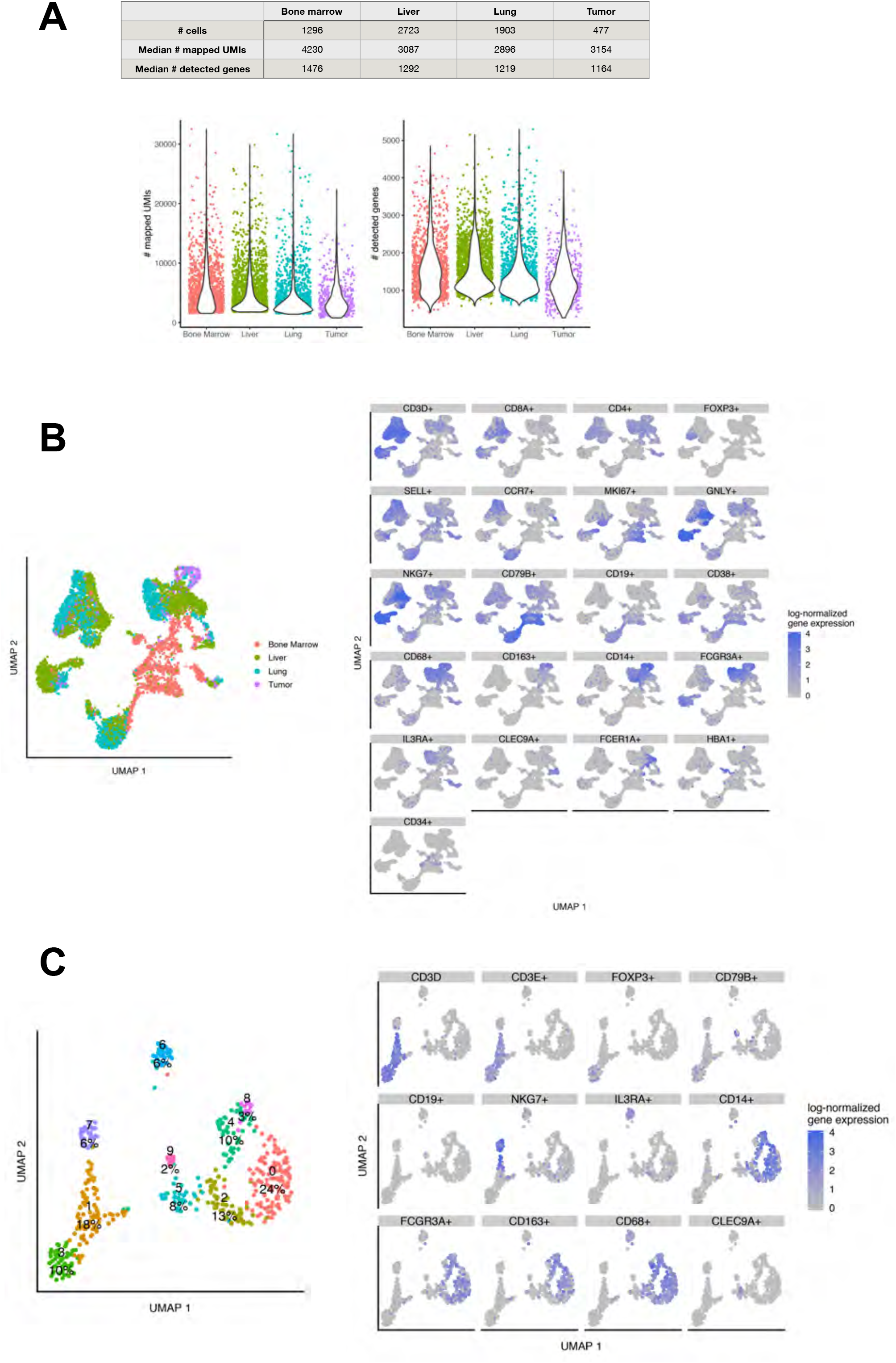
(**A**) Quality control of the scRNA-seq dataset. Distribution of the number of UMIs across tissues (bone marrow, liver, lung and tumor). (**B** and **C**) UMAP plots representing the expression of cell type-defining genes among human CD45+ cells isolated from all tissues (**B**) and from the Me275 melanoma TME (**C**) of humanized MISTRG mice.

**Supplementary Figure 2.**
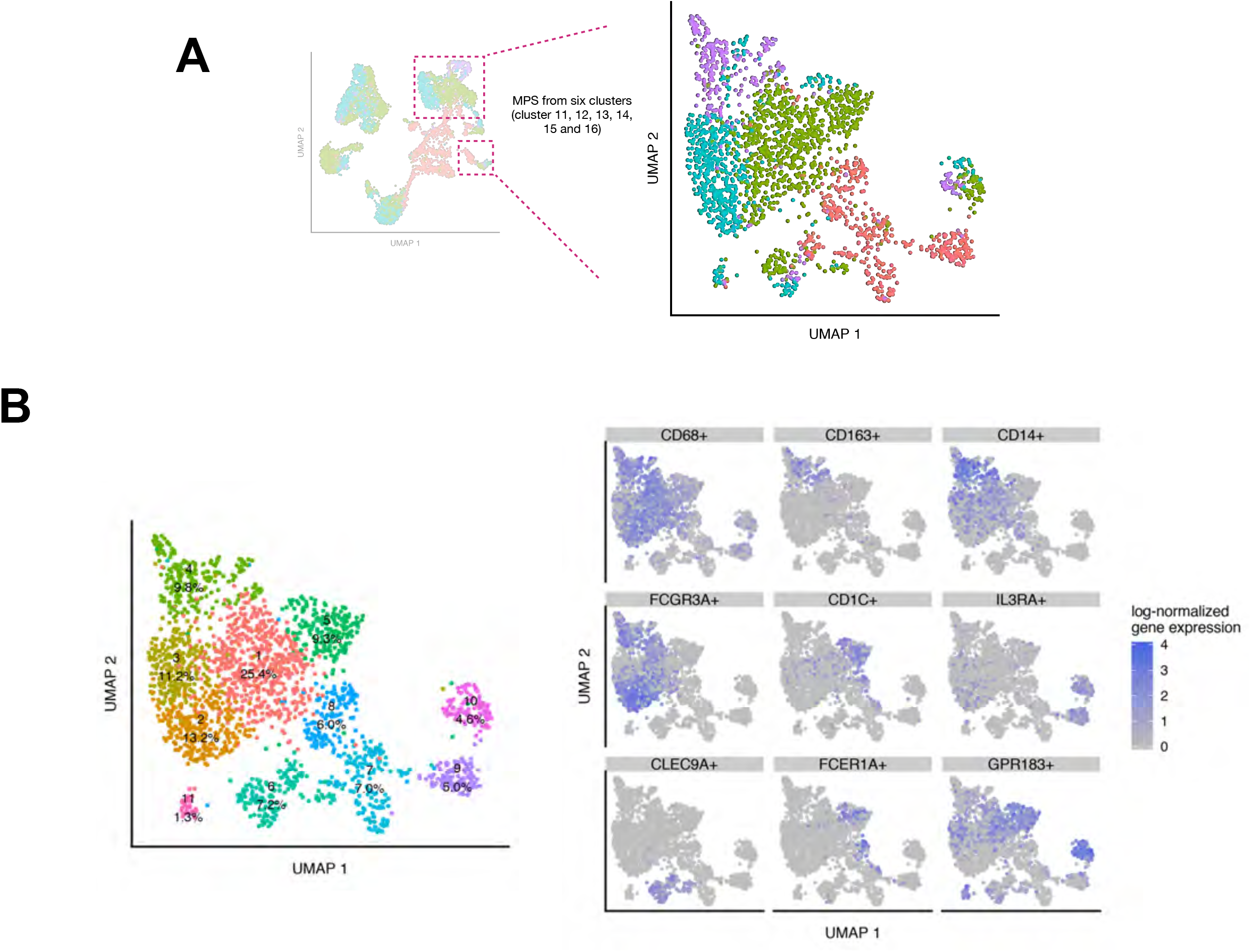
(**A**) Schematic representation of the selection of MPS cell and re-clustering. (**B**) UMAP embeddings representing the expression of cell subset-defining genes among MPS cells from all tissues.

**Supplementary Figure 3.**
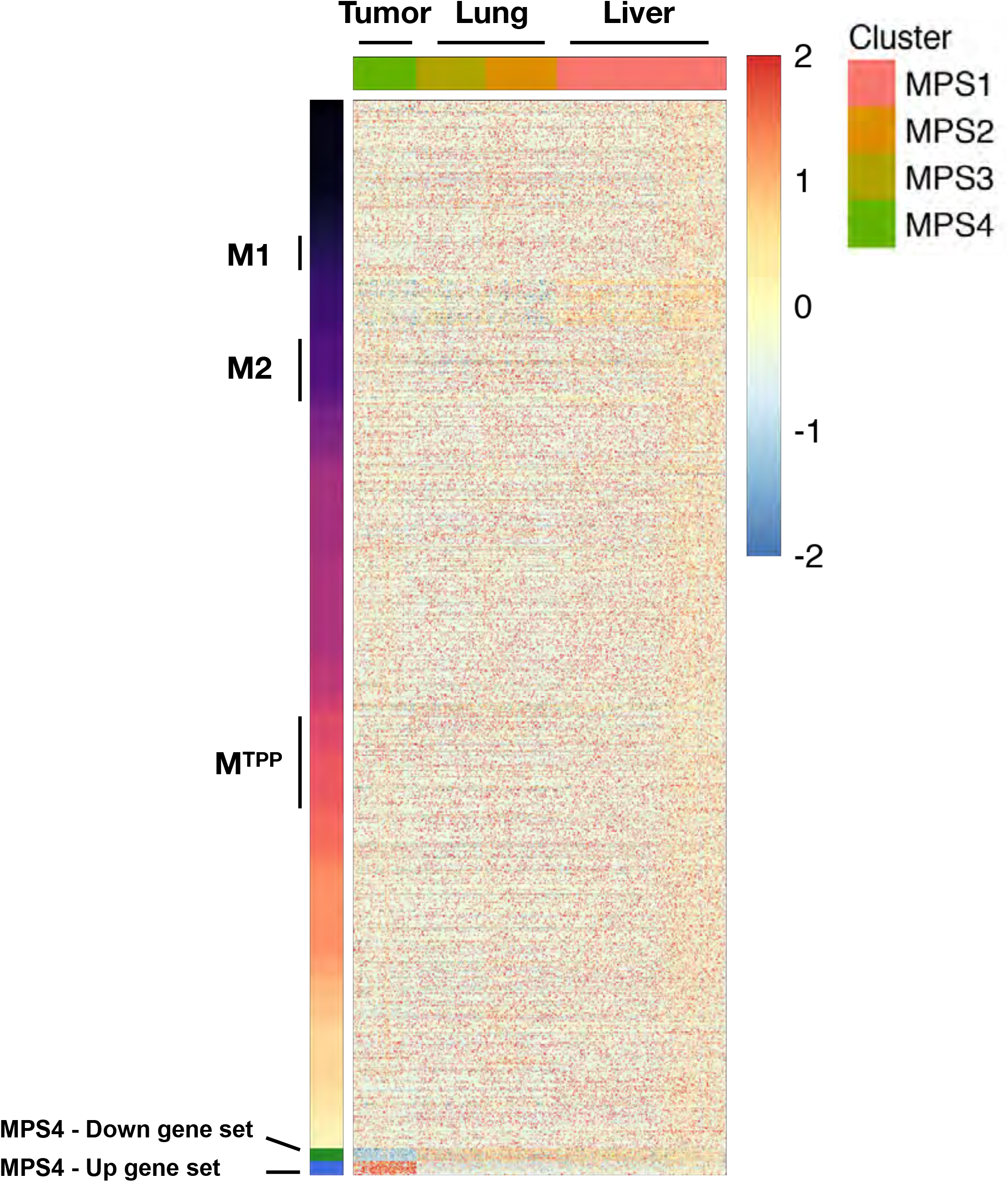
(**A**) Heatmap representing the expression, at single cell resolution of all the genes composing the 49 transcriptional modules of human macrophage polarization [32] and the MPS4 up and down signatures (Suppl. Table 2B). (**B**) Expression patterns of additional markers of immaturity and maturity along MPS pseudotime, as in Figure 4D.

**Supplementary Figure 4.**
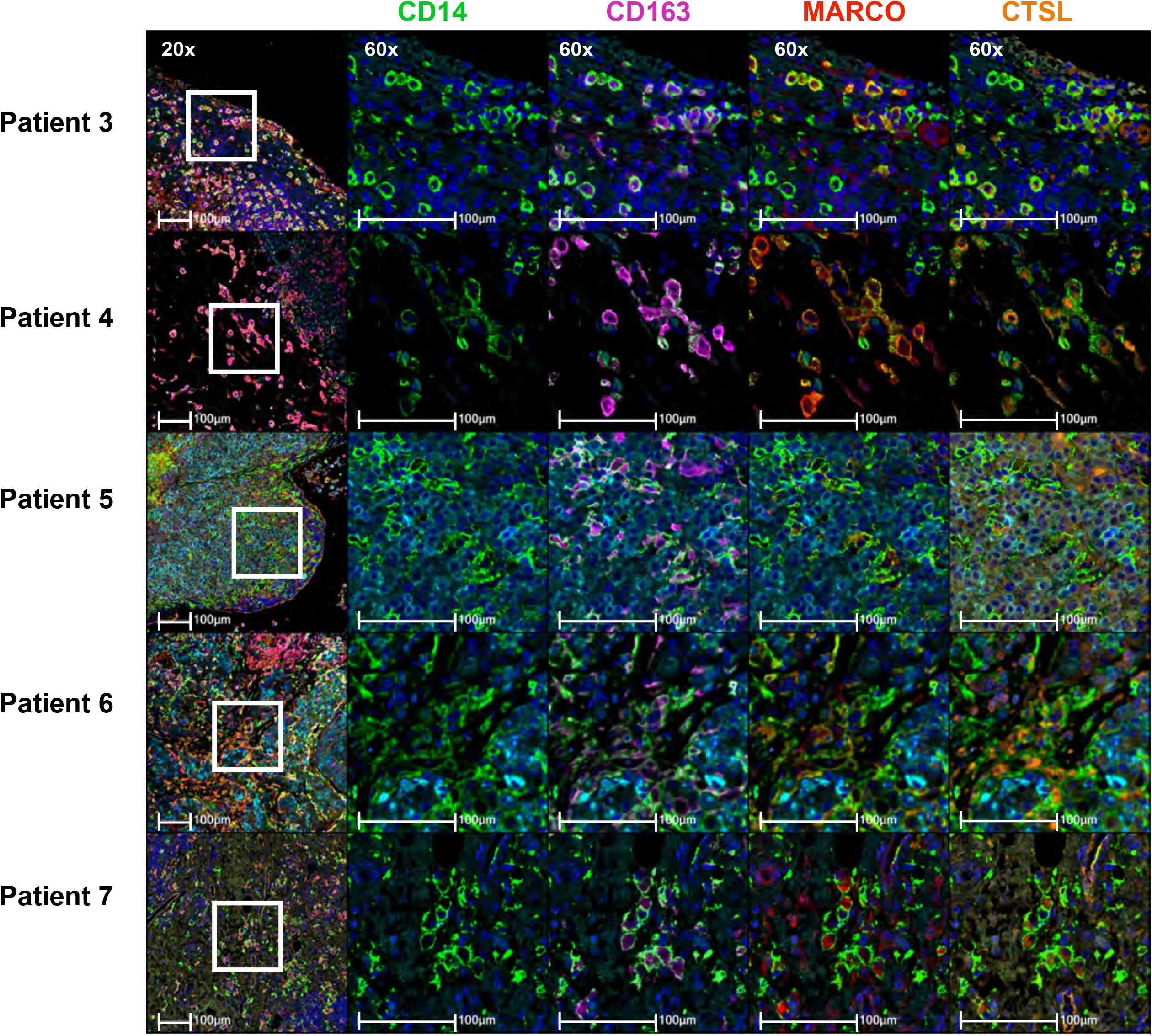
Representative images of the co-expression of MPS4 cell markers detected by multiplexed IHC in the TME of five melanoma patients (melanoma, cyan; CD14, green; CD163, magenta; MARCO, red, CTSL, orange).

**Supplementary Figure 5.**
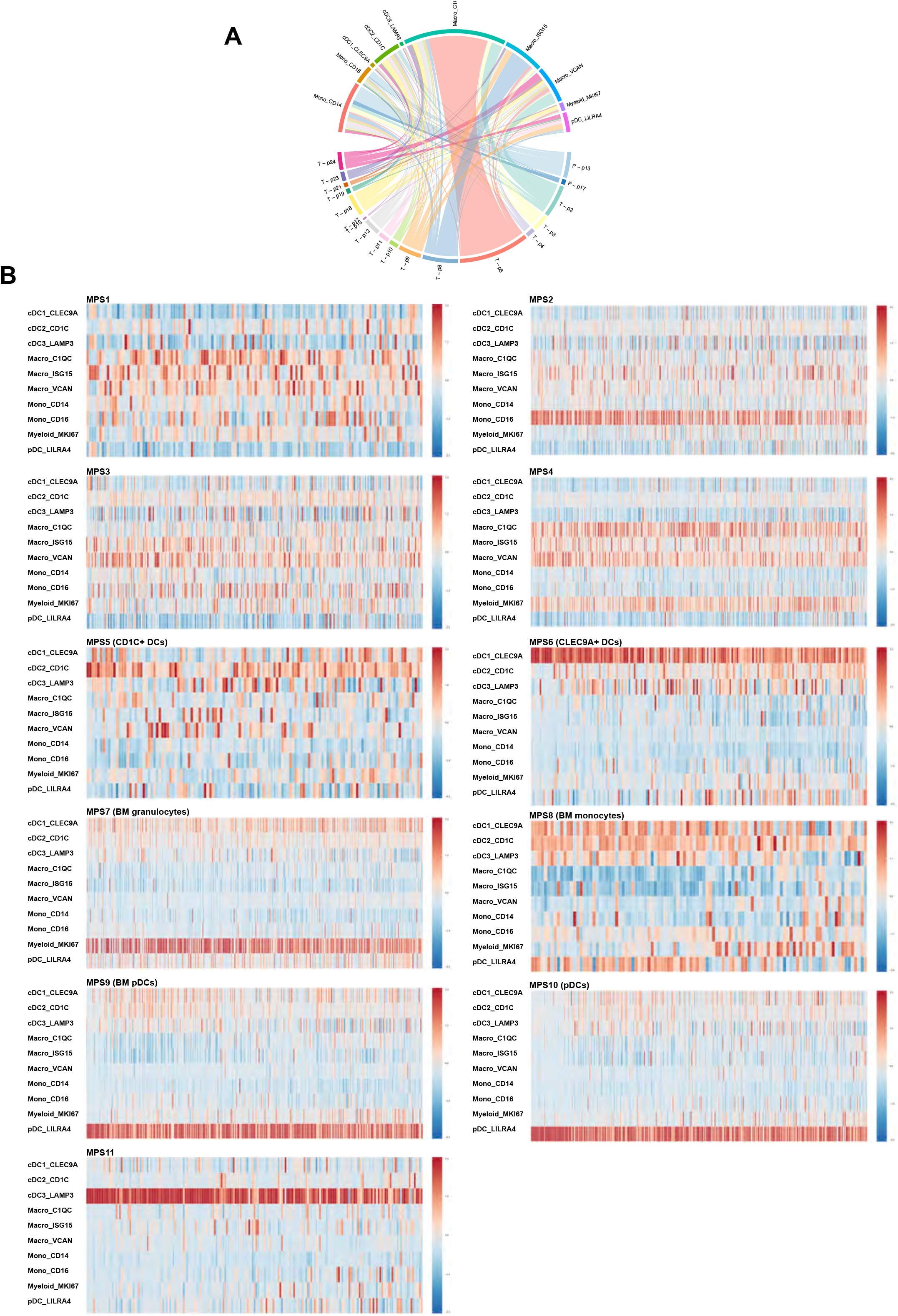
(**A**) Distribution of myeloid cell clusters, as identified by Cheng et al [16], among tumor samples (T) and PBMC (P) from melanoma patients. Data from Li et al [52], retrieved through http://panmyeloid.cancer-pku.cn/. (**B**) Expression of the MPS1-11 transcriptional signatures (from Suppl. Table 2A) by myeloid cell clusters from human melanoma patients.

**Supplementary Figure 6.**
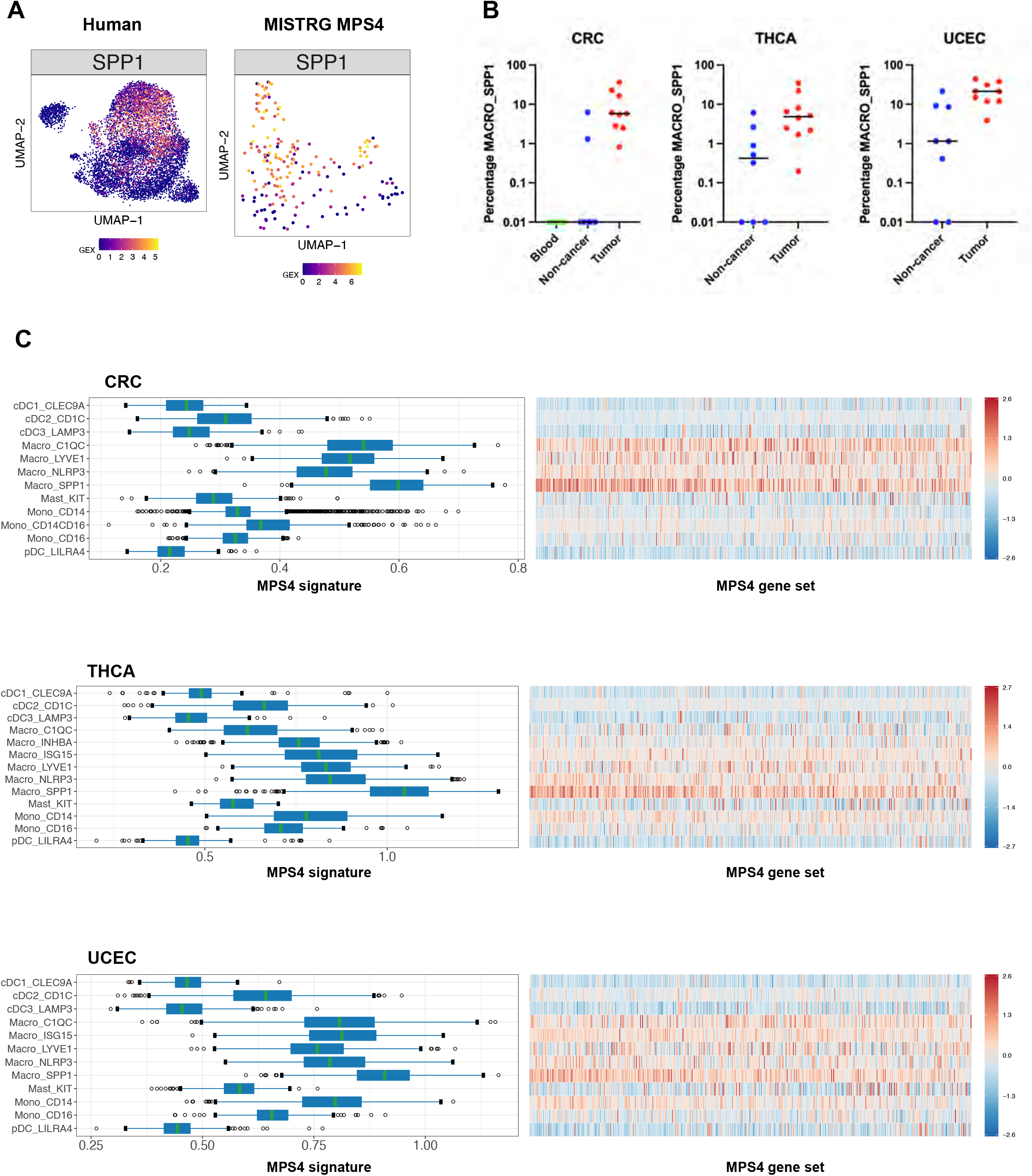
(**A**) UMAP embeddings representing the expression of *SPP1* among myeloid cells from melanoma patients, and MPS4 cells from the melanoma TME of humanized MISTRG mice. (**B**) Frequency of Macro_SPP1 cells among myeloid cells from tumors or control tissues in patient samples of three cancer types (colorectal cancer, CRC; thyroid carcinoma, THCA; uterine corpus endometrial carcinoma, UCEC). Data from Cheng et al [16], retrieved through http://panmyeloid.cancer-pku.cn/. (**C**) Expression of the MPS4 transcriptional signatures (from Suppl. Table 2A) by myeloid cell clusters from CRC, THCA and UCEC patient samples.

